# A Comprehensive Molecular Portrait of Human Urine-derived Renal Progenitor Cells

**DOI:** 10.1101/602417

**Authors:** Md Shaifur Rahman, Wasco Wruck, Lucas-Sebastian Spitzhorn, Martina Bohndorf, Soraia Martins, Fatima Asar, Audrey Ncube, Lars Erichsen, Nina Graffmann, James Adjaye

## Abstract

**Background:** Human urine is now recognised as a non-invasive source of stem cells with regeneration potential. These cells are mesenchymal stem cells but their detailed molecular and cellular identities are poorly defined. Furthermore, unlike the mouse, the gene regulatory network driving self-renewal and differentiation into functional renal cells *in vitro* remain unresolved.

**Methods:** We isolated urine stem cells from 10 individuals from both genders and distinct ages, characterized them as renal progenitor cells and explored the gene regulatory network sustaining self-renewal.

**Results:** These cells express pluripotency-associated proteins-TRA-1-60, TRA-1-81, SSEA4, C-KIT and CD133. Expression of pluripotency-associated proteins enabled rapid reprogramming into iPSCs using episomal-based plasmids without pathway perturbations. Transcriptome analysis revealed expression of a plethora of nephrogenesis-related genes such as *SIX2, OSR1, CITED1, NPHS2, NPHS1, PAX2, SALL1, AQP2, EYA1, SLC12A1* and *UMOD*. As expected, the cells transport Albumin by endocytosis. Based on this, we refer to these cells as urine derived renal progenitor cells-UdRPCs. Associated GO-term analysis of UdRPCs and UdRPC-iPSCs underlined their renal identity and functionality. Upon differentiation by WNT activation using the GSK3β-inhibitor (CHIR99021), transcriptome and KEGG pathway analysis revealed upregulation of WNT-associated genes-*AXIN2, JUN and NKD1.* Protein interaction network identified JUN- a downstream target of the WNT pathway in association with STAT3, ATF2 and MAPK1 as a putative regulator of self-renewal and differentiation in UdRPCs. Furthermore, like pluripotent stem cells, self-renewal is maintained by FGF2-driven TGFβ-SMAD2/3 pathway.

**Conclusion:** This *in vitro* model and the data presented should lay the foundation for studying nephrogenesis in man.

**Significance Statement:** Human urine is a non-invasive source of stem cells with regeneration potential. Here, we investigated the cellular and molecular identities, and the gene regulation driving self-renewal and differentiation of these cells *in vitro*. These cells express pluripotency-associated markers enabling easy reprogramming. Based on the expression of renal associated genes, proteins and functionality, we refer to these cells as urine derived renal progenitor cells-UdRPCs. CHIR99021-induced differentiation of UdRPCs activated WNT-related genes-*AXIN2, JUN and NKD1.* Protein interaction network identified JUN as a putative regulator of differentiation whereas self-renewal is maintained by FGF2-driven TGFβ-SMAD2/3. Our data will enhance understanding of the molecular identities of UdRPCs, and enable the generation of renal disease models *in vitro* and eventually kidney-associated regenerative therapies.

## Introduction

Approximately 2000 to 7000 cells are flushed out of the renal system in our urine, which contain cells of epithelial origin, erythrocytes, leukocytes, neutrophils, lymphocytes and a rare population renal stem cells.^1^ Urine stem cells which originate from the metanephric mesenchyme (MM) and those from the glomeruli are capable of giving rise to podocytes, proximal tubular cells and distal cells. These progenitor cells express renal markers such as PAX2, PAX8,^2^ SYNPO, NPHS1, PODXL and NPHS2.^3^ Interestingly, these cells exhibit stem cell properties, i.e. expression of pluripotency-associated markers such as TRA-1-60, TRA-1-81, SSEA4, C-KIT (CD117), CD133 and SSEA4; and possess high proliferation capacity as they show telomerase activity. Further, they endow multi-differentiation potential and like bone marrow derived mesenchymal stem cells express Vimentin, CD105, CD90, CD73 and not the hematopoietic stem cell markers-CD14, CD31, CD34 and CD45.^4,5^ Studies in mice have shown that Osr1, Six2, Wnt and Wt1 are required to maintain renal progenitor cells during kidney organogenesis.^6,7,8,9,10,11^ Additionally, signalling pathways such as Wnt, Fgf, Tgfβ and Notch play major roles in renal stem cell maintenance and differentiation.^12,13,14,15^

The transcription factor, Odd-skipped related 1 (Osr1), is an early marker specific for the intermediate mesenchyme (IM); knockout mice lack renal structures due to the failure to form the IM.^16^ The homeodomain transcriptional regulator Six2 is expressed in the cap mesenchyme (CM) originating from MM. Six2 positive populations can generate all cell types of the main body of the nephron.^17^ Inactivation of Six2 results in premature and ectopic renal vesicles, leading to a reduced number of nephrons and to renal hypoplasia.^18^ Mechanistically, Osr1 plays a crucial role in Six2-dependent maintenance of mouse nephron progenitors by antagonizing Wnt-directed differentiation, whereas Wt1 maintains self-renewal by modulating Fgf signals.^9,10^ Furthermore, it has been demonstrated in mice that Bmp7 promotes proliferation of nephron progenitor cells via a Jnk-dependent mechanism involving phosphorylation of Jun and Atf2.^19^

To date, research related to transcriptional regulatory control of mammalian nephrogenesis has been limited to the mouse ^6,12^ or to transcriptome “snapshots” in human.^20^ A recent study demonstrated conserved and divergent genes associated with human and mouse kidney organogenesis,^21^ thus further highlighting the need for primary human renal stem cell models to better dissect nephrogenesis at the molecular level. Furthermore, species differences need to be considered, for example, mammalian nephrons arise from a limited nephron progenitor pool through a reiterative inductive process extending over days (mouse) or weeks (human) of kidney development.^22^ Human kidney development initiates around 4 weeks of gestation and ends around 34-37 weeks of gestation. At the anatomical level, human and mouse kidney development differ in timing, scale, and global features such as lobe formation and progenitor niche organization.^21,22,23^ These are all further evidence in support of the need of a reliable and robust human renal cell culture model.

Expression of pluripotency-associated proteins has enabled rapid reprogramming of urine derived mesenchymal and epithelial cells into induced pluripotent stem cells (iPSCs).^24,25,26,27,28^ Differentiation protocols for generating kidney-associated cell types from human pluripotent stem cells have mimicked normal kidney development.^14,29,30,31^ For example, WNT activation using a GSK3β inhibitor (CHIR99021), FGF9, Activin A, Retinoic acid (RA) and BMP7 as instructive signals have been employed to derive functional podocytes, proximal renal tubules, and glomeruli.^15,32,33,34,35,36^ Despite these efforts and achievements, there will always be variabilities between differentiation protocols, the maturation state of the differentiated renal cells and genes associated with temporal maturation during human kidney organoids formation from human iPSCs.^37,38^ We propose that using native renal stem cells isolated directly from urine will circumvent most of the shortfalls and deficiencies associated with human pluripotent stem cell-based models.

Here we provide for the first time the full characterisation of UdRPCs at the transcriptome, secretome and cellular level, which has led to the identification of a gene regulatory network and associated signalling pathways that maintain their self-renewal. We anticipate that our data will enhance our meagre understanding of the properties of UdRPCs, and enable the generation of renal disease models *in vitro* and eventually kidney-associated regenerative therapies.

## Concise Methods

### Ethics

Ethical approval (Number: 5704) was granted by the ethical committee of the medical faculty of Heinrich Heine University, Düsseldorf, Germany.

### UdRPCs isolation and culture

Urine samples were collected from 10 healthy donors with diverse age, gender and ethnicity (Table 1). Isolation and expansion of the UdRPCs followed the previously established protocols.^24,28^ Adult kidney biopsy derived primary human renal epithelial cells (hREPCs) (C-12665, Promo Cell) were used as control. Description of the other cell line used in this study and culture condition has been provided in Supplemental Material and Methods.

**Table 1.**
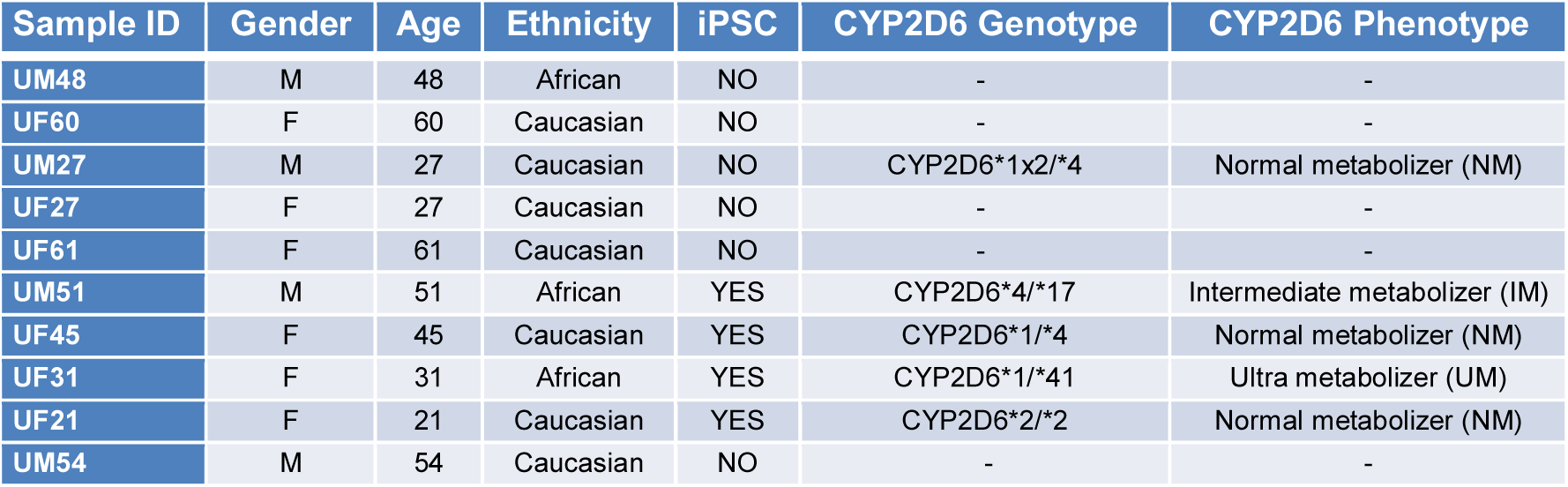
UdRPCs sample overview.

| Sample ID | Gender | Age | Ethnicity | iPSC | CYP2D6 Genotype | CYP2D6 Phenotype |
| --- | --- | --- | --- | --- | --- | --- |
| UM48 | M | 48 | African | NO | - | - |
| UF60 | F | 60 | Caucasian | NO | - | - |
| UM27 | M | 27 | Caucasian | NO | CYP2D6*1x2/*4 | Normal metabolizer (NM) |
| UF27 | F | 27 | Caucasian | NO | - | - |
| UF61 | F | 61 | Caucasian | NO | - | - |
| UM51 | M | 51 | African | YES | CYP2D6*4/*17 | Intermediate metabolizer (IM) |
| UF45 | F | 45 | Caucasian | YES | CYP2D6*1/*4 | Normal metabolizer (NM) |
| UF31 | F | 31 | African | YES | CYP2D6*1/*41 | Ultra metabolizer (UM) |
| UF21 | F | 21 | Caucasian | YES | CYP2D6*2/*2 | Normal metabolizer (NM) |
| UM54 | M | 54 | Caucasian | NO | - | - |

### CYP2D6 genotyping and phenotyping

CYP2D6 genotyping and phenotyping of five individuals were carried out by CeGat GmbH Germany using genomic DNA. The CYP2D6 variant assay reveals the pharmacogenetics (PGx) profile of an individual’s genotype and phenotype based on tested pharmacogenetics markers. The assay identifies and discriminates individuals with poor, normal, intermediate and ultra-rapid metabolizing activity.^39^

### Analysis of cell proliferation

Cell proliferation were analysed using resazurin metabolic colorimetric assay is described in details in the Supplemental Material and Methods.

### Immunophenotyping by flow cytometry

The analysis of MSC-associated cell surface marker expression of UdRPCs was performed using MSC Phenotyping Kit (Miltenyi) according to the manufactureŕs instructions and as described before.^40^ Description of the details methods has been provided in Supplemental Material and Methods.

### Albumin endocytosis assay

Albumin endocytosis assay was performed as described before. ^41^ Description of the details methods has been provided in Supplemental Material and Methods.

### Differentiation into adipocytes, chondrocytes and osteoblasts

Differentiation of UdRPCs into adipocytes, chondrocytes and osteoblasts were tested using the StemPro Adipogenesis, Chondrogenesis, and Osteogenesis differentiation Kits (Gibco, Life Technologies, CA, USA) as described before.^40,41^ A light microscope was used for imaging. Methods described in details in the Supplemental Material and Methods.

### Bisulfite genomic sequencing

Bisulfite sequencing was performed following bisulfite conversion with the EpiTec Kit (Qiagen, Hilden, Germany). Primers were designed after excluding pseudogenes or other closely related genomic sequences which could interfere with specific amplification by amplicon and primer sequences comparison in BLAT sequence database (https://genome.ucsc.edu/FAQ/FAQblat.html). In brief, the amplification conditions were denaturation at 95°C for 13min. followed by 37 cycles of 95°C for 50s, TM for 45s and 72°C for 30s. The amplification product is 469 bp in size. Amplification product was cloned into pCR2.1vector using the TA Cloning Kit (Invitrogen, Carlsbad, United States) according to the manufacturer’s instructions. On average 30 clones were sequenced using the BigDye Terminator Cycle Sequencing Kit (Applied Biosystems, Foster City, United States) on a DNA analyzer 3700 (Applied Biosystems) with M13 primer to obtain a representative methylation profile the OCT4 promoter region. 5’-regulatory gene sequences are defined by +1 transcription start of the following sequence: Homo sapiens POU class 5 homeobox 1 (POU5F1), transcript variant 1, mRNA. NCBI Reference Sequence: NM_002701.6

### Generation of iPSC from UdRPCs

UdRPCs were reprogrammed into iPSCs using an integration-free episomal based transfection system without pathway inhibition. Briefly, UdRPCs were nucleofected with two plasmids pEP4 E02S ET2K (Addgene plasmid #20927) and pEP4 E02S CK2M EN2L (Addgene plasmid #20924) expressing a combination of pluripotency factors including OCT4, SOX2, LIN28, c-MYC, KLF4, and NANOG using the Amaxa 4D-Nucleofector Kit according to the manufacturer’s guidelines and as described previously.^28^

### Immunofluorescence study and Western blot analysis

Immunofluorescence study was performed as described previously.^40^ List of primary antibody presents in Supplementary Table S1. Details methods of Immunofluorescence study and Western blot analysis have been described in Supplemental Material and Methods.

### Quantitative RT-PCR analysis

Real-time quantitative PCR was performed in technical triplicates with Power SYBR Green Master Mix (Life Technologies), 12.5 ng cDNA per sample and 0.6 μM primers on a VIIA7 (Life Technologies) machine. Mean values were normalized to levels of the housekeeping gene ribosomal protein L37A calculated by the 2−ΔΔCt method. Primers used were purchased from MWG (primer sequences and predicted sizes of amplicons presented in supplementary table S2).

### Culture supernatant analysis

For the detection of cytokines secreted by the UdRPCs, we employed the Proteome Profiler Human Cytokine Array Panel A (R&D Systems, MA, USA) following the manufacturer’s instructions. Correlation variations and *p* values were calculated based on the pixel density.

### Microarray data analyses

Total RNA (1μg) preparations were hybridized on the PrimeView Human Gene Expression Array (Affymetrix, Thermo Fisher Scientific) at the core facility Biomedizinisches Forschungszentrum (BMFZ) of the Heinrich Heine University Düsseldorf. Gene expression data will be available online at the National Center of Biotechnology Information (NCBI) Gene Expression Omnibus. The raw data was imported into the R/Bioconductor environment ^42^ and further processed with the package affy ^43^ using background-correction, logarithmic (base 2) transformation and normalization with the Robust Multi-array Average (RMA) method. The heatmap.2 function from the gplots package ^44^ was applied for cluster analysis and to generate heatmaps using Pearson correlation as similarity measure. Gene expression was detected as previously described ^45^ using a detection-p-value threshold of 0.05. Differential gene expression was determined via the p-value from the limma package ^46^ which was adjusted for false discovery rate using the q value package.^47^ Thresholds of 1.33 and 0.75 were used for up-/down-regulation of ratios and 0.05 for p-values. Venn diagrams were generated with the VennDiagram package.^48^ Subsets from the venn diagrams were used for follow-up GO and pathway analyses.

### KEGG pathway, GO and network analysis

Gene ontology (GOs) terms were analysed within the Bioconductor environment employing the package GOstats.^49^ GOs of category Biological Process (BP) were further summarized with the REVIGO tool ^50^ to generate treemaps populating the parameter for allowed similarity with tiny=0.4. GO networks were generated from the REVIGO tool in xgmml format and imported into Cytoscape.^51^ To reduce the network to a readable size they were filtered in Cytoscape by the log10(p) between −3.75 and −2.75. The saturation of the red nodes representing GO terms indicates the significance via the p-value while the grey value of the edges represents their similarity. KEGG pathways ^52^ were downloaded from the KEGG server in March 2018 and tested for over-representation with the R-built-in hypergeometric test.

### Activated WNT pathway associated protein interaction network

The network was constructed from the 20 most significantly up- and down down-regulated genes between CHIR99021 treatment and untreated controls. Genes were ranked by the limma-p-value and passed the criteria: detection p-value < 0.05 for the dedicated condition, ratio < 0.75 or ratio > 1.33, limma-p-value < 0.05. The resulting 40 genes are marked as green nodes in the network. Interacting proteins containing at least one protein coded by the 40 genes were retrieved from BioGrid version 3.4.161.^53^ To reduce complexity and increase visualization, the network was minimized by adding only the n=30 interacting proteins (marked as red nodes) with the most interactions to proteins coded by the 40 genes. The plot of the interactions network was drawn employing the R package network.^54^ Communities of related proteins within the network were detected employing an in-betweenness clustering analysis via the method cluster_edge_betweenness () from the R package igraph.^55^

### Statistics

All data are presented as arithmetic means ± standard error of mean. At least 3 independent experiments were used for the calculation of mean values. Statistical analysis was performed by U-test and student’s t-test. P values of <0.05 were considered significant.

## Results

### UdRPCs express a subset of pluripotent stem cell-associated markers and possess features typical of bone marrow-derived MSC

Urine samples were collected from 10 healthy adult donors (4 males-UM and 6 females-UF) with ages ranging from 21 to 61 years, and of mixed ethnicity (3 african and 7 caucasian) (Table 1). Attached cells emerged as isolated clusters after 7 days, thereafter these acquired a “rice grain” fibroblast-like morphology resembling MSCs (Figure 1A). A selection of distinct UdRPC populations (n=4) were used to assay cell proliferation and growth. After 3 days in culture, the cells exited the lag phase and growth began in an exponential phase. Cells attained stationary phase at day 7 of subculture, and entered in a decline phase after 7-9 days (Figure 1B). All four populations-UM27, UM16, UM51 and UF45 showed similar proliferation and growth patterns.

**Figure 1.**
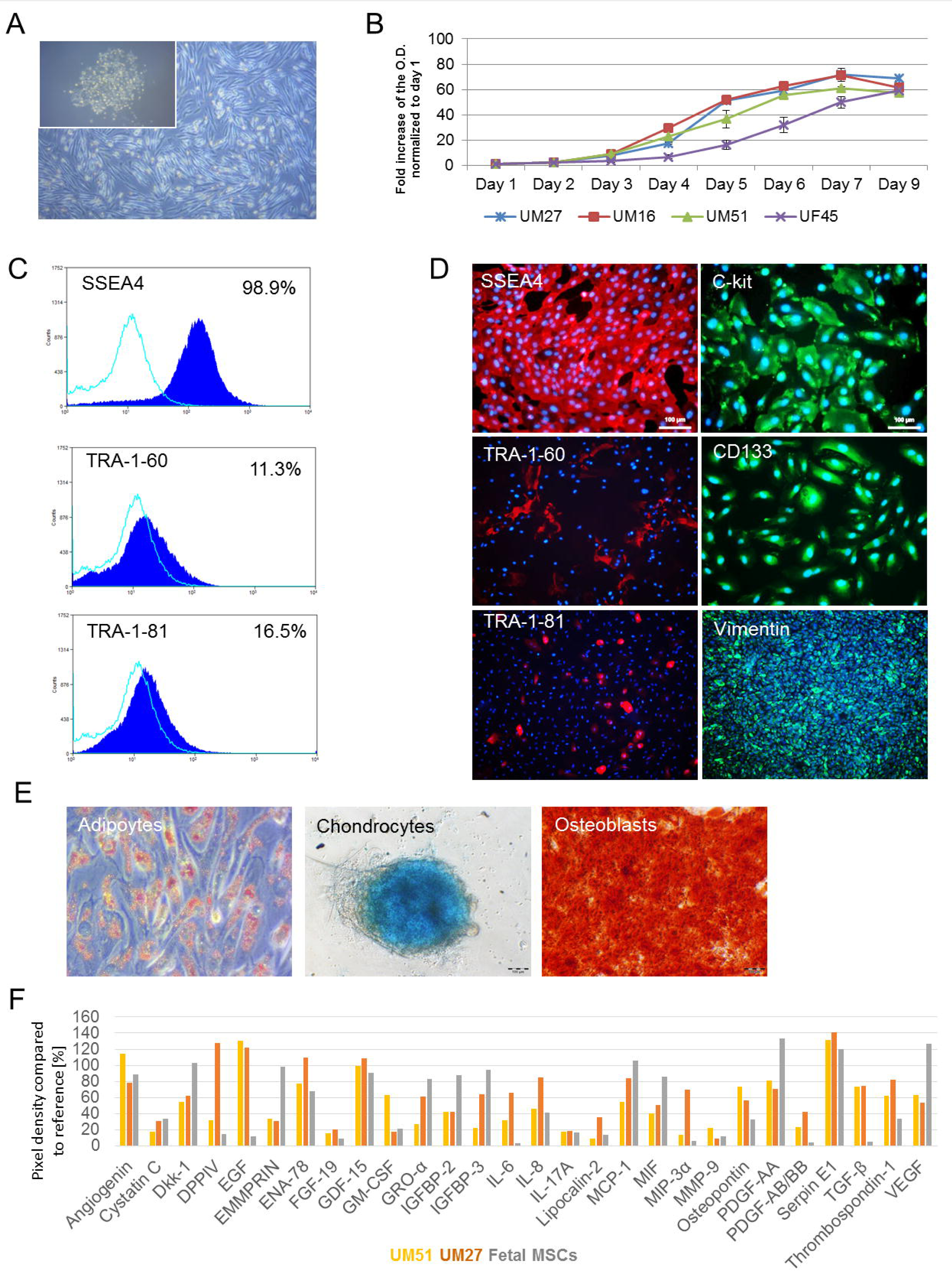
Propagation and characterisation of UdRPCs. (A) Representative pictures of the “rice grain”-like appearance of the cells from the initial attachment to an elongated MSC-like morphology. (B) Growth curve analysis of selected UdRPCs carried out using the Resazurin metabolic assay. Data are presented as means ±SEMs. (C) Immune-phenotyping and (D) immunofluorescence-based detection of the expression of pluripotency-associated stem cell-proteins (SSEA4 (red), TRA-1-60 (red), TRA-1-81 (red), C-KIT (green), CD133 (green) and the mesenchymal-associated protein Vimentin (green); cell nuclei were stained using Hoechst/DAPI (scale bars: 100µm). (E) *In vitro* Osteoblast, Chondrocyte and Adipocyte differentiation potential of UdRPCs. (F) Cytokines secreted by UdRPCs in culture media. Lists of significant GÒs and KEGG pathways associated with the genes encoding the secreted cytokines, are shown in Supplemental Figure S1.

Flow cytometry analysis revealed that approximately 98.9% of the cells express SSEA4, TRA-1-60 (11.3%) and TRA-1-81 (16.5%) (Figure 1C). These data were confirmed by immunofluorescent-based staining which also showed expression of the typical mesenchymal marker-Vimentin and the proliferation-associated stem cell markers-C-KIT and CD133 (Figure 1D). In order to reveal the detailed methylation pattern of the 5’-regulatory region of the OCT4 gene in iPSCs derived from the UM51 and the corresponding parental cells (control) we employed standard bisulfite sequencing. In total 330 Cytosine-phosphatidyl-Guanine-dinucleotides (CpG) slightly upstream of the transcription starting site (TSS) of the OCT4 gene were analysed. Within this 469bp long region, a dense methylation pattern was observed in the UM51 control cells, with 92.4% (305) of the CpG dinucleotides identified were methylated. In contrast, iPSCs derived from UM51 had 72.12% (207) of analysed CpGs were unmethylated (Supplemental Figure S1).

Flow cytometry analysis of critical MSC cell surface markers were negative for the hematopoietic markers CD14, CD20, CD34, and CD45 and positive for CD73, CD90 and CD105 albeit at variable levels (Supplemental Figure Figure S2). Typical of MSCs, UdRPCs can be driven to differentiate into osteocytes, chondrocytes, and adipocytes when cultured in the respective differentiation medium for 3 weeks (Figure 1E). Furthermore, employing a cytokine array (n=2), a repertoire of trophic factors such as IL8, GDF-15, SERPINE-1, Angiogenin, VEGF, and Thrombospondin-1 was detected as secreted in a similar manner as fetal-femur derived MSCs (Figure 1F, Supplemental Figure S2).

The CYP2D6 genotypes investigated were distinct between groups of individuals, thus reflecting the diverse drug metabolizing activity between individuals. UM51 for example expresses the CYP2D6 *4/*17 genotype which confers an intermediate metabolizing activity^28^ whereas UF31 bears the CYP2D6*1/*41 genotype which confers an ultra-rapid metabolizing activity. The other three individuals (UF21, UF45 and UM27) are endowed with normal drug metabolizing rates (Table 1).

### UdRPCs express key renal progenitor cell markers and are able to endocytose Albumin

Immunofluorescence-based staining revealed expression of the key proteins including transcription factors associated with kidney development-SIX2, CITED1, WT1, Nephrin, LHX1, and PAX8 as shown by representative pictures (Figure 2A). Additionally they were shown to express Cytokeratin 19 and PODXL (Supplemental Figure S2) and transport Albumin (Figure 2B).

**Figure 2.**
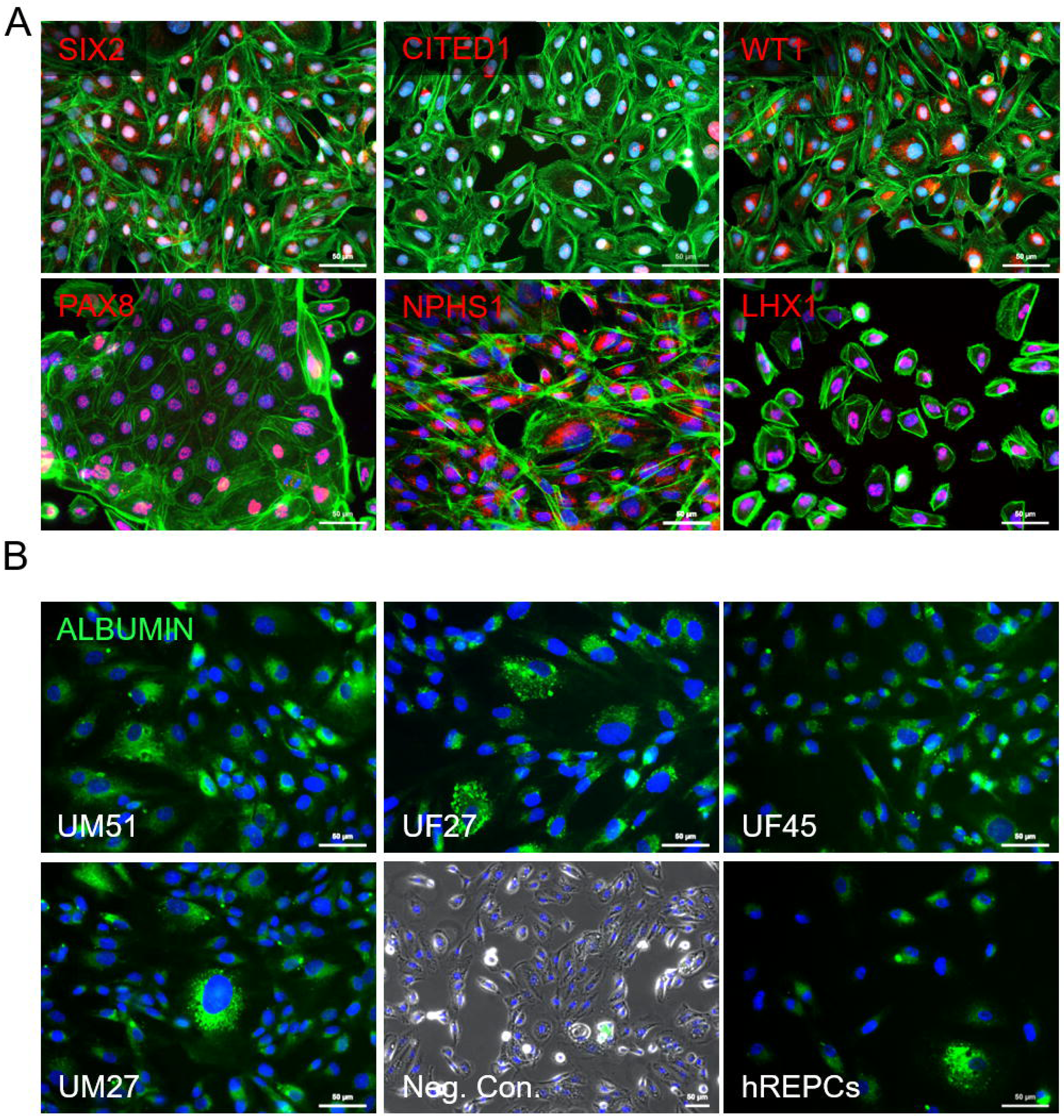
Expression of kidney-associated proteins in UdRPCs and Albumin transport. (A) UdRPCs express the renal markers-SIX2, CITED1, WT1, PAX8, Nephrin (NPHS1) and LHX1. Renal markers (red), phalloidin (green), cell nuclei were stained using DAPI/Hoechst (blue). (B) UdRPCs (n=4) like the human kidney biopsy-derived hREPCs also transport Albumin. Albumin was coupled to Alexa Fluor 488 (green) and cell nuclei stained with DAPI (blue). Scale bars indicate 50µm.

### Comparative transcriptome analysis of UdRPCs and kidney biopsy derived hREPCs

A hierarchical clustering and heatmap analysis was carried out to compare the transcriptomes of UdRPCs with the kidney biopsy-derived renal epithelial proximal cells (hREPCs). As anticipated, all UdRPCs clustered together as a common cell type distinct from hREPCs.

UdRPCs express higher levels of renal progenitor cell markers such as *SIX2*, *CITED1*, *UMOD*, *PAX2*, *NPHS2*, *GDNF*, *SALL4*, *MIXL1* and *OSR1*. This heatmap also revealed that UdRPCs are of mesenchymal origin expressing *VIM* and *SIX2* whereas the hREPCs are differentiated epithelial expressing *CDH1* and elevated levels of *GATA3* and *SOX17* (Figure 3A). Direct comparison of logarithmic (base 2) gene expression values of UdRPCs, for instance UM51 with hREPCs, in a scatter plot (Figure 3B) shows similarity with a high Pearson correlation of 0.9575. The more epithelial character of hREPCs is reflected by *CDH1* which has one of the highest ratios between hREPCs and UM51 r = 156.96. Additionally, an elevated expression of the mesenchymal marker *VIM* in UM51 r = 1.1 confirmed the mesenchymal phenotype of UdRPCs. The comparison of expressed genes (det-p < 0.05) in UdRPCs (UM51) and hREPCs in a venn diagram (Figure 3C) showed that most genes are expressed in common (12281), whereas 566 are expressed exclusively in UM51 and 438 exclusively in hREPCs. The 10 most over-represented GO BP terms (biological processes) in the UM51 exclusive gene-set include triglyceride homeostasis, kidney development and urogenital system development, whereas the hREPCs exclusive gene set includes chloride transmembrane transport, anion transport and response to lipopolysaccharides (Figure 3D). The common gene set consists of 874 up-regulated genes (ratio > 2) in UM51 (e.g. renal tubule development, urogenital system development and anterior/posterior pattern specification) and 1042 down-regulated genes (ratio < 0.5) in UM51 (e.g. cell division and cholesterol biosynthetic process) (Figure 3E).

**Figure 3.**
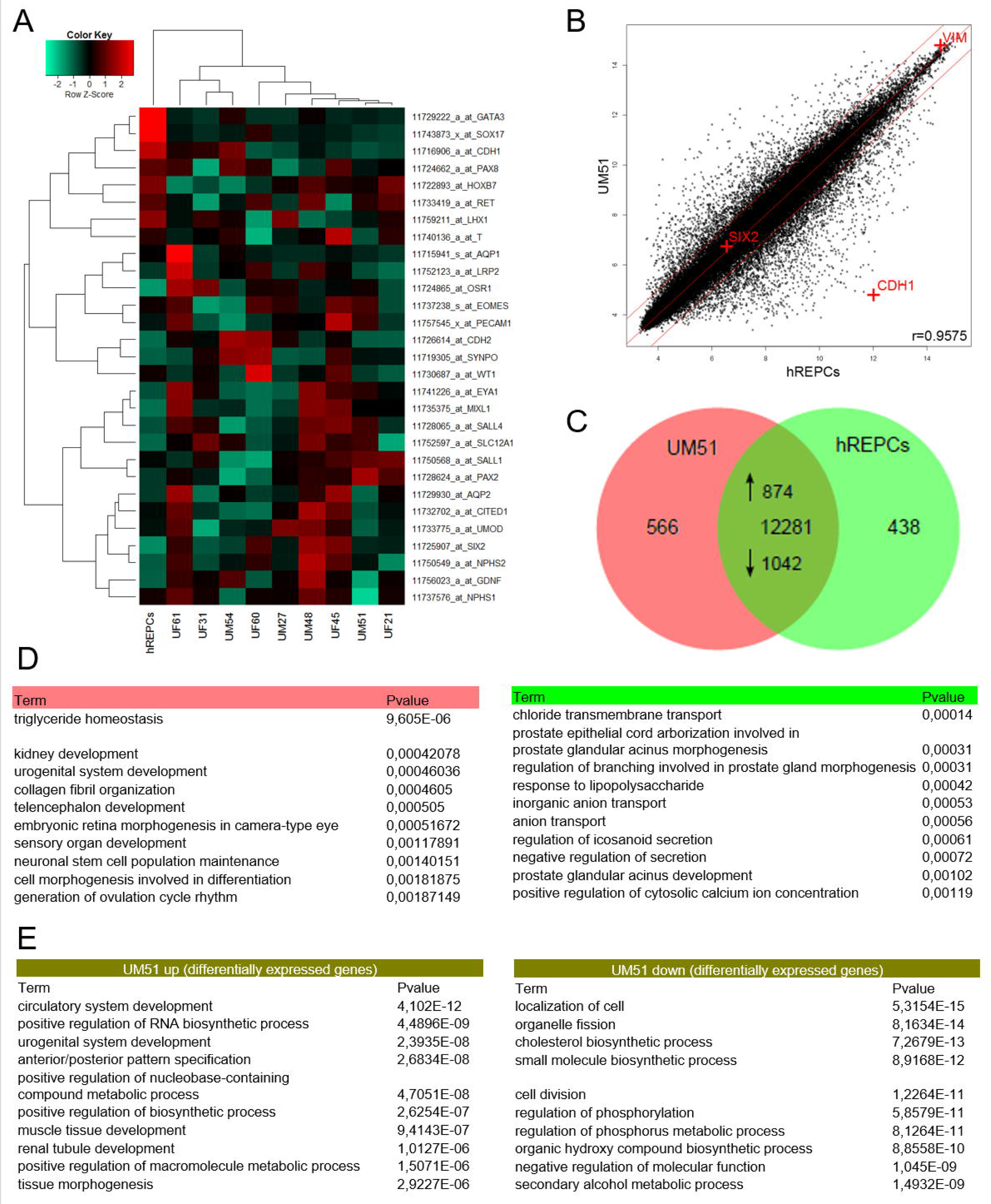
Transcriptome analysis of UdRPCs in comparison to kidney biopsy-derived renal epithelial proximal cells-hREPCs. (A) The heatmap of kidney-specific markers expressed in UdRPCs and hREPCs. (B) Comparison of gene expression values of UdRPCs (UM51) with hREPCs in a scatter plot confirms the mesenchymal phenotype of UdRPCs, i.e. expression of Vimentin (*VIM*) and expression of E-cadherin (*CDH1*) in hREPCs (C) Expressed genes (det-p < 0.05) in UdRPCs (sample UM51) and hREPCs are compared in the venn diagram. (D) The 10 most over-represented GO BP-terms in 566 UM51 genes include triglyceride homeostasis and kidney development and in 438 hREPCs genes include chloride transmembrane transport. (E) The 10 most over-represented GO BP-terms in the up- and down-regulated genes in UM51 in comparison to hREPCs are shown. The complete dataset is presented in Supplemental Table S3.

### Confirmation of the renal origin of UdRPCs and retention of renal-associated genes in UdRPC derived iPSCs

A venn diagram-based comparison of gene expression (det-p < 0.05) in UdRPCs and human foreskin fibroblasts (HFF) was carried out (Figure 4A) in order to dissect common and distinct gene expression patterns. The majority of genes (11649) are expressed in common, 463 exclusively in UdRPCs and 891 in fibroblasts. The 463 genes were further analysed for over-represented GOs and summarized as a GO network (Figure 4B) with the tools REVIGO, and Cytoscape was used for the GO terms of the category BP. In addition to several developmental terms such as organ induction, regulation of embryonic development (high number of edges referring to similarity to many terms), specific renal-related terms including urogenital system development, mesenchymal cell proliferation involved in ureteric bud development and positive regulation of nephron tubule epithelial cell differentiation (marked with blue ellipse, intense red indicating higher significance) were identified. Interestingly, the non-canonical Wnt signalling pathway, which plays a major role in kidney development, is also over-represented (orange ring-top left).

**Figure 4.**
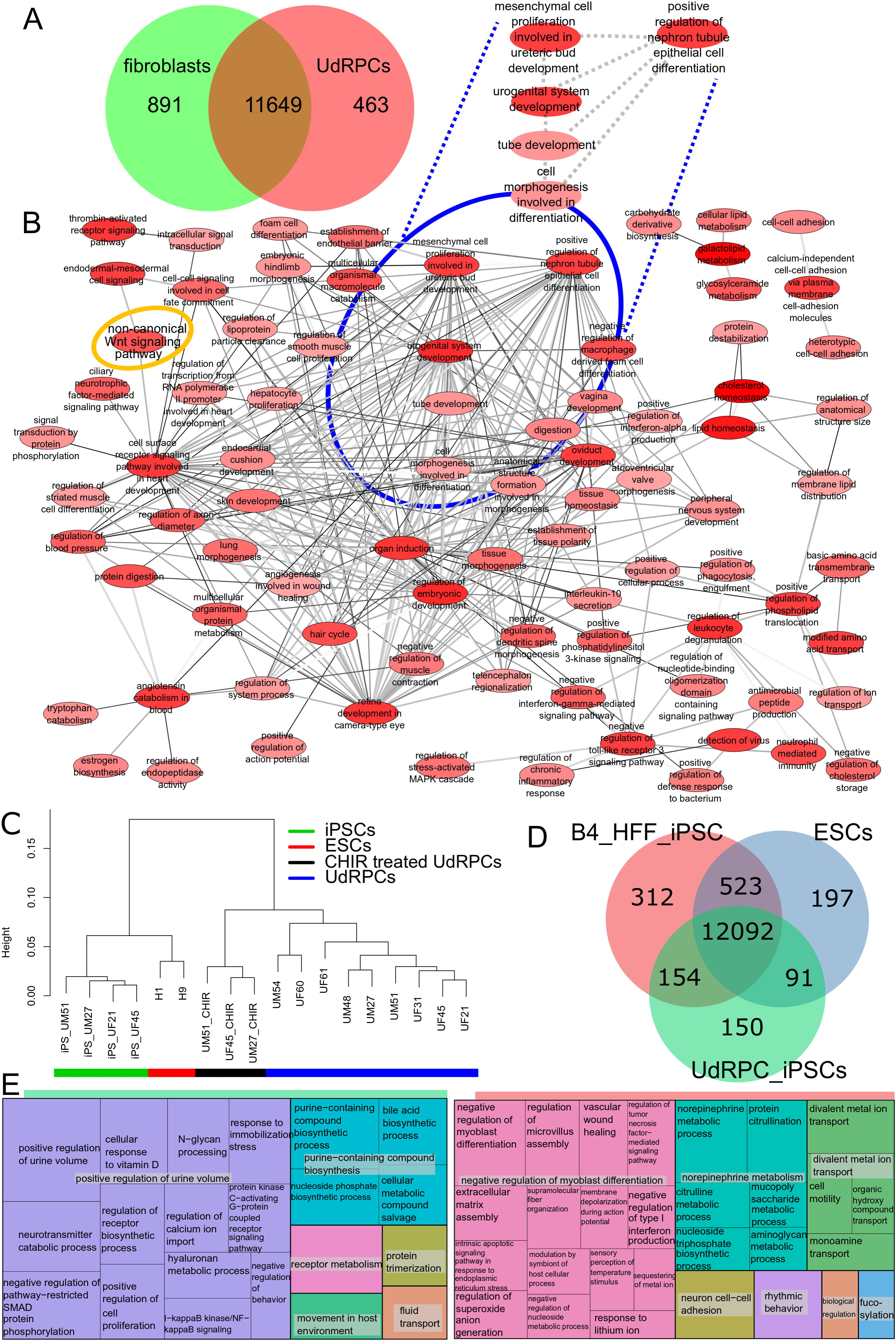
In-depth bioinformatic analysis of UdRPCs and UdRPC-iPSCs. (A) Expressed genes (det-p < 0.05) in UdRPCs and fibroblasts are compared in a venn diagram. Most genes are expressed in common (11649), 463 genes are expressed exclusively in UdRPCs and 891 in fibroblasts. The subsets and UdRPCs GOs are presented in supplemental_table_S4. (B) The gene ontology network was generated with the tools REVIGO and Cytoscape and summarizes the GO terms of category Biological Process (BP) over-represented in the 463 genes expressed exclusively in UdRPCs. Several general developmental terms emerged, e.g. “organ induction”. Specific renal-related terms including “urogenital system development” are marked with a blue ellipse. GOs are represented by network nodes with the intensity of red indicating the significance of over-representation of a GO term. The edges refer to similarities between the GO terms. (C) The dendrogram shows a clear separation of UdRPCs, differentiated UdRPCs (black bar), ESCs (H1 and H9, red bar) and UdRPCs-iPSCs (green bar). (D) Venn diagram of HFF-iPSCs, UdRPCs-iPSCs and ESCs. (E) GO terms of 150 genes expressed exclusively in UdRPC-iPSCs indicate that UdRPC-iPSCs retain the memory of renal origin. In the treemap for the HFF-iPSCs the GO-BP terms of the 312 over-represented genes of the exclusive gene set are summarized. The most significant group is associated with negative regulation of myoblast differentiation including genes *DDIT3*, *MBNL3*, *TGFB1*, *ZFHX3* pointing at the fibroblast origin of these iPSCs. The entire dataset is presented in Supplemental Table S5.

The dendrogram based on the global transcriptome analysis revealed a clear separation of UdRPCs lines (n=9) from the differentiated UdRPCs (CHIR 99021 treated UdRPCs, n=3), UdRPCs-iPSCs (n=4) and embryonic stem cells (H1 and H9) (Figure 4C). Characterization of the derived UdRPC-iPSCs is depicted in Supplemental Figure S2 (Supplemental Figure S3A, S3B, S3C, and S3D). In the venn diagram (Figure 4D) we compared expressed genes (det-p < 0.05) in UdRPC-iPSCs with ESCs and HFF-iPSCs. Most genes (12092) are expressed in common in all cell types while 150 genes are expressed exclusively in UdRPC-iPSCs. The genes expressed exclusively in one cell type were further analysed for over-representation of GO terms. The treemap summarizing the GO terms of category BP over-represented in the 150 genes expressed exclusively in UdRPC-iPSCs (Figure 4E) indicates that UdRPC-iPSCs retain a memory of their kidney origin. In addition to the largest most significant group-positive regulation of urine volume, it consists of other renal-related GO terms (e.g. calcium transport, vitamin D). Stem-cell-related and developmental terms such as positive regulation of cell proliferation are due to their pluripotent nature. Within the treemap summarizing the GO-BP terms over-represented in the 312 genes expressed exclusively in HFF-iPSCs, the largest most significant group is associated with negative regulation of myoblast differentiation, thus pointing at the fibroblast origin of these iPSCs (Figure 4E). Furthermore, within the treemap summarizing the GO-BP terms over-represented in the 197 genes expressed exclusively in ESCs, the largest most significant group is associated with negative regulation of astrocyte differentiation-hinting at their known propensity to differentiate into the ectodermal lineage (Supplemental Figure S4).

### WNT pathway activation by GSK3**β** inhibition induces differentiation of UdRPCs into renal epithelial proximal tubular cells

To differentiate 3 independent UdRPC preparations, the cells were treated with 10 μM CHIR99021 (WNT pathway activation by GSK3β inhibition) for 2 days and morphological changes from fibroblastic to elongated tubular shape were observed (Figure 5A). In the venn diagram, expressed genes (det-p < 0.05) in untreated UdRPCs are compared to UdRPCs treated with CHIR99021. Genes expressed in common amounts to 11790, of these 2491 are upregulated in the CHIR99021 treatment (p < 0.05, ratio > 1.33) and 2043 are down-regulated (p < 0.05, ratio < 0.75) (Figure 5B, Supplemental Table S6). Among the upregulated genes, 27 are considered “novel” (gene symbol starting with “LOC”), 21 among the down-regulated genes and 98 among the non-regulated genes (Supplemental Table S6). The heatmap based on the top 20 regulated genes shows a clear separation between untreated and treated cells (Figure 5C). Amongst the up-regulated genes, the associated KEGG pathways include WNT-signaling (*AXIN2*, *JUN*, *NKD1*) (Supplemental Figure S5). Over-representation analysis of the up-regulated genes and their associated KEGG pathways identified protein processing in endoplasmic reticulum as highly significant and several signalling pathways such as mTOR, Insulin, p53, AMPK and TNF. Over-representation analysis of the down-regulated genes and associated KEGG pathways revealed cell cycle, cellular senescence, focal adhesion, FoxO, ErbB and thyroid hormone signalling. Interestingly Hippo pathway was regulated in both undifferentiated and differentiated UdRPCs (Figure 5D).

**Figure 5.**
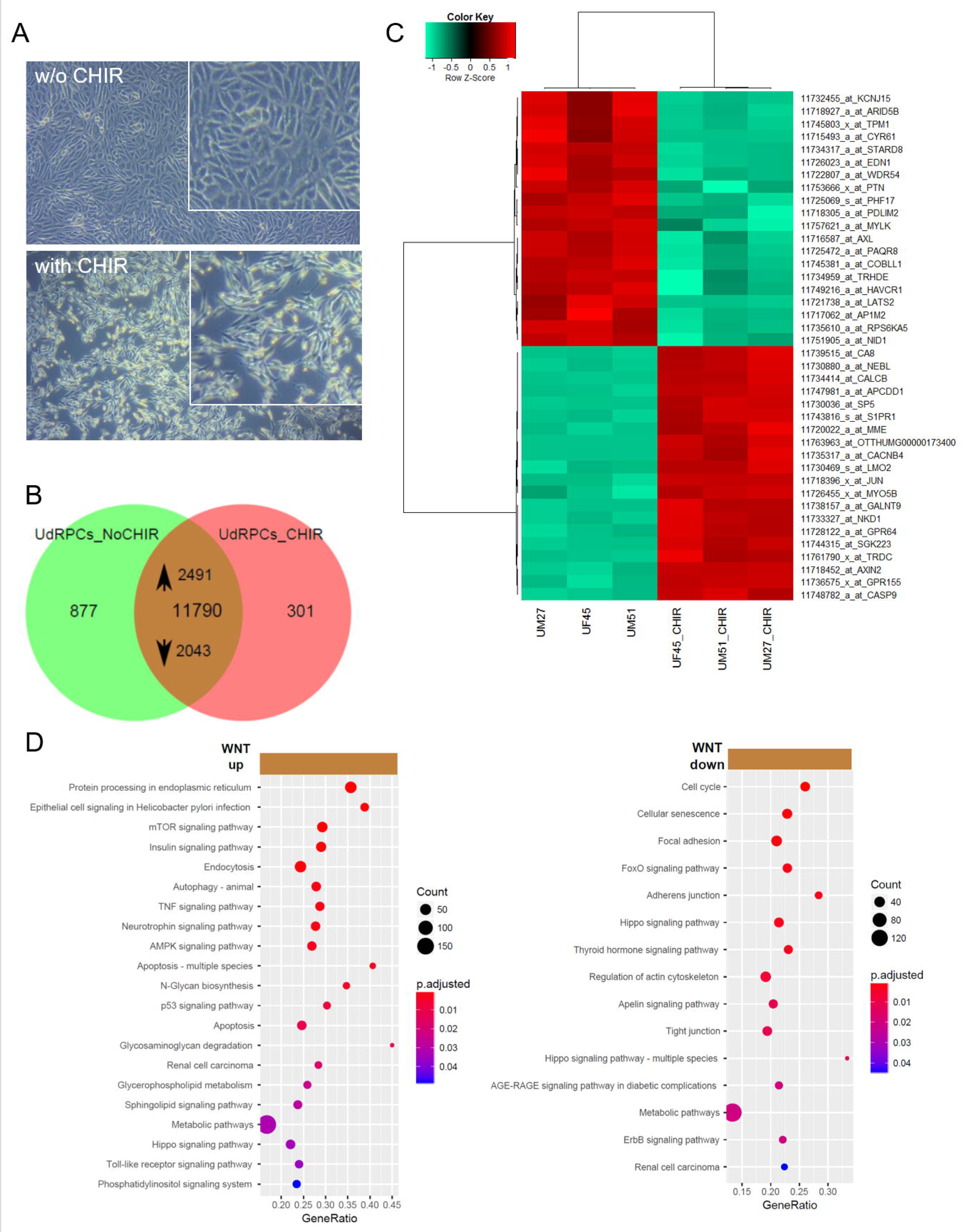
Supplementation of UdRPCs with the GSK-3β inhibitor. (A) Activation of WNT signalling by supplementation with GSK-3β-inhibitor CHIR99021 led to differentiation into renal epithelial proximal tubular cells. (B) Heatmap of 3 independent UdRPC preparations with and without CHIR treatment. (C) In the venn diagram, expressed genes (det-p < 0.05) in untreated UdRPCs are compared to UdRPCs treated with the GSK-3β-inhibitor CHIR99021. Among the 11790 genes expressed in both conditions, 2491 are up-regulated in the CHIR99021 treatment (p < 0.05, ratio > 1.33) and 2043 down-regulated (p < 0.05, ratio < 0.75). (D) Over-representation analysis of the up-regulated genes and associated KEGG pathways revealed protein processing in endoplasmic reticulum as highly significant and several signalling and metabolic pathways including mTOR, Insulin, p53 and TNF. Over-representation analysis of the down-regulated genes in KEGG pathways identified cell cycle, cellular senescence, focal adhesion, FoxO and adherens junction as most significant. Supplemental Table S6 provides the full list of regulated genes and associated pathways.

### Regulation of self-renewal and differentiation in UdRPCs

Further to the transcriptome analyses (Figure 5), Real-time PCR revealed downregulation of the stem cell self-renewal associated gene *CD133* and activated expression of the nephrogenesis-associated gene *BMP7* after CHIR stimulation (Figure 6A).

**Figure 6.**
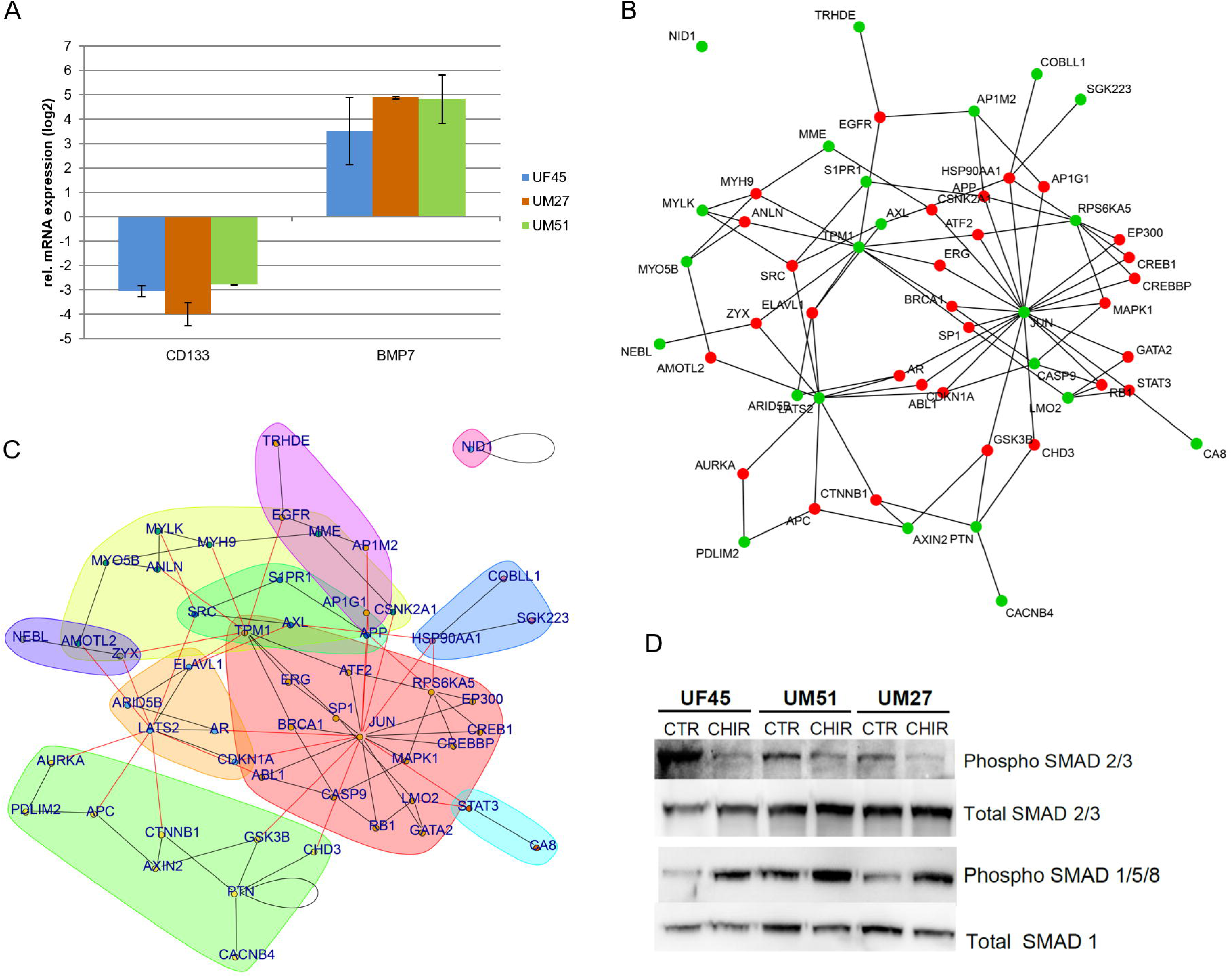
Regulation of self-renewal and differentiation in UdRPCs. (A) Real-time PCR-based confirmation of down-regulation of *CD133* and activated expression of *BMP7* after CHIR stimulation. (B) JUN is a major hub of protein interaction networks of UdRPCs treated with CHIR. Based on the Biogrid database protein interaction networks were constructed from the set of the most highly regulated 40 genes either up- or down in the UdRPCs treated with CHIR. The selected genes used to connect to the network with interactions from the Biogrid database are marked in green, genes added as Biogrid interactions are marked in red. Induction of WNT leading to GSK3B inhibition is reflected by the connection of GSK3B to JUN and to AXIN2 which is connected to CTNNB1 (β-catenin) – these all downstream targets of GSK3B in the WNT-signaling pathway. (C) Community clustering of the network identified several communities: JUN (red), GSK3B/AXIN2/CTNNB1 (green), LATS2 (yellow), EGFR (pink). Black lines refer to edges within a community, red lines to edges between different communities. (D) Western blot analysis of the phosphorylated levels of SMAD 2/3 and SMAD 1/5/8 in undifferentiated and differentiated UF45, UM51 and UM27.

To identify key self-renewal regulators and pathways in UdRPCs, a protein-protein-interaction network was generated. The network of the 40 proteins, encoded by the 20 most significantly up- and down-regulated genes between CHIR treated and untreated UdRPCs (Figure 5C) indentified JUN as a major hub – in terms of having most connections to other proteins in the network. However, in the WNT-signaling pathway JUN is at the end of a downstream cascade from GSK3B, including further downstream targets-AXIN2 and CTNNB1. The genes encoding these proteins were differentially regulated by the CHIR treatment (green nodes) (Figure 6B). Several communities with more interactions within the community than to other communities can be detected in the network via community clustering of the network via edge-betweenness includes JUN (red), GSK3B / AXIN2 / CTNNB1 (green), LATS2 (yellow), EGFR (pink) (Figure 6C). To analyze the effect of WNT activation on the TGFβ-SMAD pathway, Western blot analysis was performed to detect phosphorylation levels of SMAD 2/3 and SMAD 1/5/8 in UF45, UM51 and UM27. In the differentiated cells (UdRPCs after CHIR treatment) a decreased level of phosphorylated SMAD 2/3 and increased levels of phosphorylated SMAD 1/5/8 were observed (Figure 6D).

## Discussion

Here we describe urine as a reliable, robust and cheap source of renal stem cells, in contrast to amniotic fluid or kidney biopsies.^41,56^ Urine can be accessed non-invasively, without risk for the patient and repeated samples can be collected from the same donor over prolonged periods. Urine derived stem cells can be expand from a single clone with high proliferation potency.^24,57^ We propose naming these cells as urine derived renal progenitor cells-UdRPCs, because they can be kept in culture for almost 12 passages whilst maintaining expression of the self-renewal associated proteins-CD133, C-KIT, TRA-1-60, TRA-1-81 and SSEA4 (Figure 1D) as has been shown by others^24,41^ Despite the expression of these pluripotency-associated factors, UdRPCs do not express OCT4, SOX2 and NANOG- which are key pluripotency-regulating transcription factors.^58,59^ Further evidence in support of the lack of OCT4 expression is our observed OCT4 promoter methylation pattern in the UM51 cells, with 92.4% (305) of the CpG dinucleotides identified as methylated whereas iPSCs derived from UM51 had 72.12% (207) of analysed CpGs unmethylated (Supplemental Figure S1).

UdRPCs express key renal progenitor-regulatory proteins SIX2, CITED1, WT1 and NPHS1 (Figure 2A) indicating they originate from the kidney as described from others.^13,56,60^ Furthermore, UdRPCs transport Albumin (Figure 2B) as observed in renal stem cells.^41,61^ We have shown that UdRPCs are in fact bon-fide MSCs- i.e. they express VIM and not CDH1, adhere to plastic surfaces, express CD73, CD90 and CD105 and not the hematopoietic markers CD14, CD20, CD34, and CD45 (Supplemental Figure S2). Typical of MSCs, UdRPCs can be differentiated into osteoblasts, chondrocytes and adipocytes (Figure 1E).^41,56,62^ They also secrete a plethora of cytokines and growth factors-such as EGF, GDF, PDGF and Serpin E1 (Figure 1F).^63^

With regards to nephrogenesis-associated regulatory genes,^17,30^ we observed *SIX2*, *CITED1*, *GDNF*, *WT1*, *NPHS1*, *PAX2* expression in UdRPCs but at variable expression levels between individual cell preparations (Figure 3A). The GOs network derived from the exclusively expressed genes in UdRPCs (compared to HFF) unveiled renal system development-related terms (Figure 4B). Furthermore, the GOs from the UdRPCs-iPSC exclusive genes set, in contrast to pluripotent stem cells, identified terms related to renal function therefore implying the preservation of their kidney origin (Figure 4E). As the conservation of tissue of origin in iPSCs might be linked to epigenetic memory,^4,64^ UdRPCs as well as UdRPCs-iPSCs, especially with known CYP2D6 status, might be advantageous for differentiation into renal cells, modelling kidney-related diseases, nephrotoxicity studies and regenerative medicine. However, dissecting the gene regulatory mechanisms that drive human renal progenitor growth and differentiation *in vitro* represents the key step for translation but remains a challenge due to the absence of well-characterised urine derived stem cells. Here we have shown that UdRPCs are a self-renewing stem cell population unlike the kidney biopsy-derived hREPCs which are differentiated renal epithelial cells (Figure 3A, Figure 3B). To demonstrate that UdRPCs can maintain self-renewal when cultured under undifferentiation conditions but yet retain the potential for epithelial differentiation and nephrogenesis, we induced active WNT signalling, by treatment with the GSK-3β inhibitor-CHIR99021. The differentiated cells adopted an elongated tubular morphology (Figure 5A) and reduced proliferation as also shown for human ESC and iPSC derived renal epithelial cells.^65,66,67^

Although WNT pathway activation induced an epithelial phenotype, we did not see a dramatic increase in *CDH1* expression but rather activation of *CDH-3* expression (8.86 fold) (Supplemental Table S6). *Cdh-3*, a gene encoding a member of the cadherin superfamily, functions in epithelial cell morphogenesis in *Caenorhabditis elegans*^68^ an event which is poorly understood in human nephrogenesis.

In line with our previously published observations in amniotic fluidic-derived renal cells, the down-regulated expression of *SIX2*, *WT1*, *CD133* and upregulated expression of *BMP7* (Figure 6A) induced the loss of self-renewal.^41^ Global transcriptome analyses also revealed the down-regulation of 2043 genes some of which are associated with pathways such as cell cycle, FoxO, Hippo and ErbB signalling (Figure 5D). The Hippo pathway which is composed of WNT target genes such as *LATS2, AXIN2* and *CTNNB1* have been reported to regulate epithelialization of nephron progenitors.^69,70^

We detected differential expression 40 genes in which 20 most significantly up- and down-regulated between WNT-induced differentiated and self-renewing UdRPCs (Figure 5C). Amongst the genes up regulated in the CHIR treated cells are the WNT targets-*AXIN2*, *JUN* and *NKD1* known to be associated with WNT signalling (Supplemental Figure S5). Interestingly, a protein interaction network identified JUN as a major hub connected to GSK3β and interlinked with ATF2, STAT3, GATA2 and MAPK1 (Figure 6B, Figure 6C). Interestingly, in a mouse nephrogenesis model, Bmp7 phosphorylates Jun and Atf2 via Jnk signalling which promote the proliferation of mouse nephron progenitors.^19^ This indeed might be contradictory to our observed elevated expression of *BMP7* upon WNT induced differentiation of UdRPCs- i.e. suppression of *BMP7* expression is needed to maintain self-renewal in UdRPCs. It is well known that TGFβ signals through pSMAD2/3 whereas BMPs activate SMAD1/5/8. Since, SMADs are a target of MAPK particularly of JNK, BMPs and TGFβ both can activate the SMAD circuit.^71,72^

Based on this study and our previously published data in human amniotic fluid-derived renal cells,^41^ we propose that similar to self-renewal in human pluripotent stem cells, UdRPCs maintain self-renewal by active FGF signalling leading to phosphorylated TGFβ-SMAD2/3 (Figure 6D).^59,73^ In contrast, activation of WNT/β-catenin signalling leads to an upregulation of *JUN* and *BMP7* leading to activation of SMAD1/5/8 signalling (Figure 6D) and exit of self-renewal by downregulation of *WT1, SIX2*, *CITED1*, and *CD133* expression. To surmise, we derived a hypothetic scheme of the WNTβ catenin and TGFβ pathway-mediated cell fate decisions in UdRPCs. This simplistic model is depicted in Figure 7.

**Figure 7.**
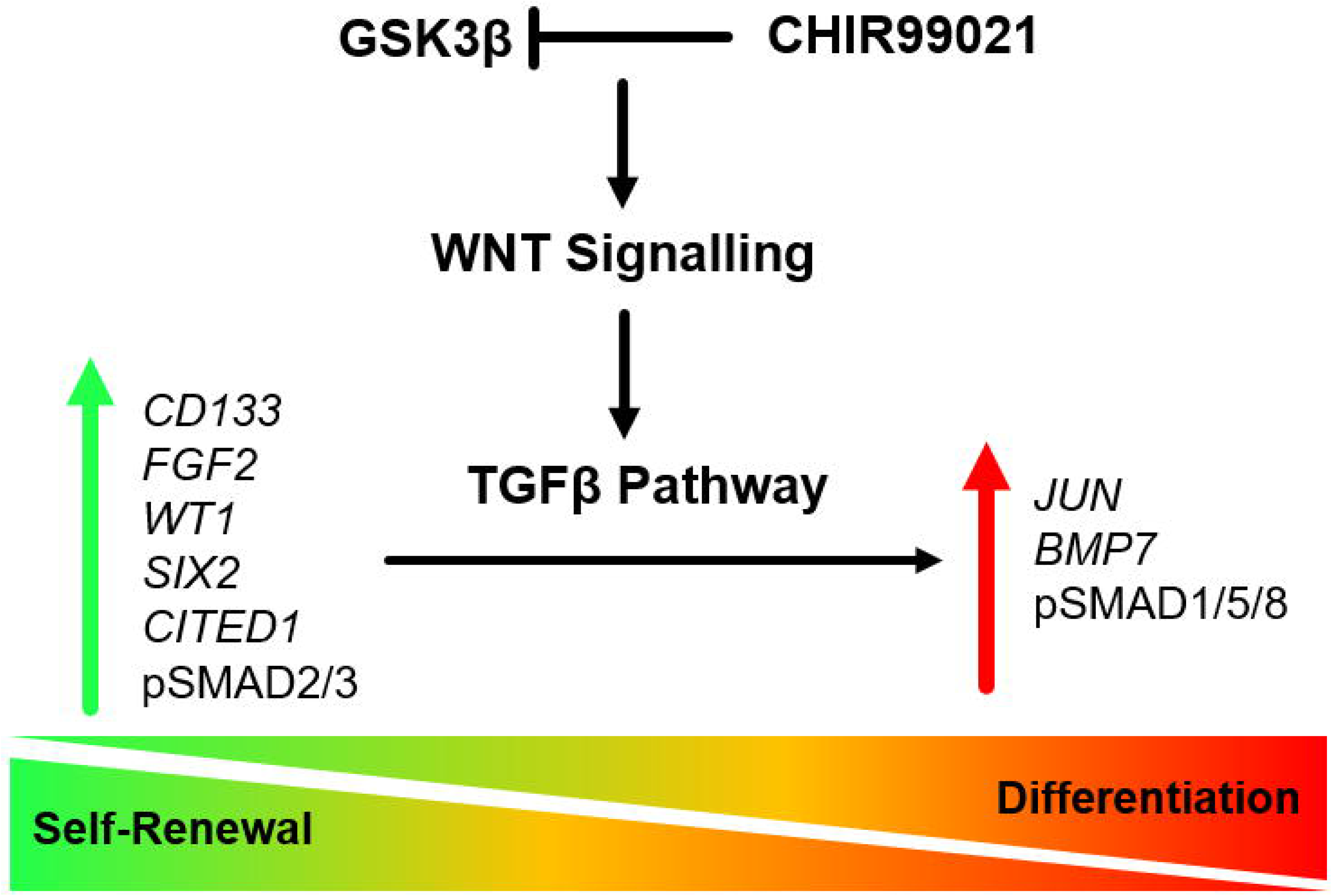
WNTβ catenin and TGFβ pathway-mediated cell fate decisions in UdRPCs. Self-renewal (inactive WNT/βcatenin signalling and active TGFβ-SMAD2/3 signalling) is maintained by elevated expression of the renal progenitor markers *SIX2*, *WT1*, *CITED1*, *CD133*, in addition to phospho-SMAD2/3 and FGF2 resulting in and down regulated expression of *BMP7*. In contrast, activation of WNT/β-catenin signalling induces upregulated expression of JUN and *BMP7* leading to activation of phospho-SMAD1/5/8, downregulated expression of *WT1, SIX2, CITED1, FGF2, CD133* and ultimately exit of self-renewal.

Comparing self-renewal of renal progenitor cells in both human (UdRPCs) and mouse, it is clear that an intricate balance is needed between SIX2, WT1, CITED1 expression and Wnt/β-catenin activity in order to determine the cell fate of nephron progenitor cells.^7,11,17,21^ Furthermore, it remains to be determined if indeed there exist subtle human and mouse differences in the gene regulatory network needed to maintain a self-renewing renal progenitor pool in both species and we believe that human UdRPCs as described here will facilitate these studies.

## Data access

All raw and processed data used in this study have been archived in NCBI gene expression omnibus under GEO accession number GSE128281. (https://www.ncbi.nlm.nih.gov/geo/query/acc.cgi?acc=GSE128281).

## Author contributions

Conception: J. A and M.B; Cell culture experiments: M.S.R, L.S.S, F.A, M.B and J.A; Real Time PCR: F.A; Bisulphite sequencing: L.E; Bioinformatics: W.W and J.A; Proliferation assays and Western blot analyses: S.M; Derivation and characterisation of iPSCs: M.B, N.G and A.N; FACS analysis: M.B; Writing of manuscript: M.S.R, W.W, L.S.S and J.A; Final edits: J.A; and all authors approved the final version of the manuscript.

## Acknowledgements

James Adjaye acknowledges support from the medical faculty of the Heinrich Heine University-Düsseldorf, Germany. Md Shaifur Rahman acknowledges support from the German Academic Exchange Service (DAAD-91607303).

## Supplemental Material and Methods

### UdRPCs differentiation

For differentiation of the UdRPCs 10µM CHIR99021 was added to the cell culture medium for 2 days. Adult kidney biopsy derived primary human renal epithelial cells (hREPCs) (C-12665, Promo Cell, Heidelberg, Germany) were used as control.

### Immunophenotyping by flow cytometry

The analysis of MSC-associated cell surface marker expression of UdRPCs was performed using MSC Phenotyping Kit (Miltenyi) according to the manufactureŕs instructions. In case of pluripotency-associated markers, TRA-1-60, TRA-1-81, and SSEA4 dye-coupled antibodies were used (anti-TRA-1-60-PE, human (clone: REA157), number 130-100-347; anti-TRA-1-81-PE, human (clone: REA246), number 130-101-410, and anti-SSEA-4-PE, human (clone: REA101), number 130-098-369; Miltenyi Biotec GmbH). Flow cytometric analysis of the stained cells was performed via BD FACSCanto (BD Biosciences, Heidelberg, Germany) and CyAn ADP (Beckman Coulter, CA, USA). Histograms were generated using the Summit 4.3.02 software.

### RNA isolation and cDNA synthesis

RNA was isolated using the Direct-zol RNA MiniPrep Kit (Zymo Research, CA, USA) according to provider guidelines. After checking the quality of mRNA, 500 ng of RNA were used for complementary DNA synthesized with the TaqMan Reverse Transcription Kit (Applied Biosystems).

### Culture supernatant analysis

For the detection of cytokines secreted by the UdRPCs, we employed the Proteome Profiler Human Cytokine Array Panel A (R&D Systems, MA, USA) following the manufacturer’s instructions. 1.5 ml of conditioned medium from cultured UdRPCs at a density of 95% was used. The array was evaluated by detection of the emitted chemiluminescence. The pixel density of each spotted cytokine was analysed using the software ImageJ. All spots on the membrane including reference and negative control spots were measured separately. Correlation variations and *p* values were calculated based on the pixel density.

### Differentiation into adipocytes, chondrocytes and osteoblasts

Differentiation of UdRPCs into adipocytes, chondrocytes and osteoblasts were tested using the StemPro Adipogenesis, Chondrogenesis, and Osteogenesis differentiation Kits (Gibco, Life Technologies, CA, USA). After the differentiation periods, cells were fixed using 4% PFA for 20 min at RT and stained with Oil Red-O for detecting adipocytes, Alcian Blue for chondrocytes, and Alizarin Red S for osteoblasts as described previously. A light microscope was used for imaging.

### Generation of iPSC from UdRPCs

UdRPCs were reprogrammed into iPSCs using an integration-free episomal based transfection system without pathway inhibition. Briefly, UdRPCs were nucleofected with two plasmids pEP4 E02S ET2K (Addgene plasmid #20927) and pEP4 E02S CK2M EN2L (Addgene plasmid #20924) expressing a combination of pluripotency factors including OCT4, SOX2, LIN28, c-MYC, KLF4, and NANOG using the Amaxa 4D-Nucleofector Kit according to the manufacturer’s guidelines and as described previously. The nucleofected cells were cultured on Matrigel coated 6-well plate containing StemMACs or mTeSR media under hypoxic conditions. Emerging colonies were picked and transferred to a new plate and cultured under normoxic conditions. After few passaging, vector-dilution PCR and genomic DNA fingerprinting were performed. Karyotyping was performed at the Institute of Human Genetics and Anthropology, Heinrich Heine University, Düsseldorf. Finally, embryoid body (EB) formation and analysis were carried out.

### Immunofluorescence-based detection of protein expression

To analyse expression of specific markers, cells were fixed with 4% PFA (Polysciences Inc., PA, USA) for 15 min at room temperature (RT) and washed three times in PBS and permeability was increased using 1% Triton X-100 for 5 min. Next, for blocking we used: 10% normal goat serum (NGS; Sigma), 0.5% Triton X-100, 1% BSA (Sigma) and 0.05% Tween 20 (Sigma) in PBS for 2h. The cells were incubated with primary antibodies (Supplementary Table S1) for 1h at RT followed by three washes with PBS. Thereafter the corresponding secondary Cy3-labeled or Alexa Fluor 488-labeled antibodies (Thermo Fisher Scientific) and Hoechst 33,258 dye (Sigma-Aldrich Chemie GmbH, Taufkirchen, Germany) or DAPI (Southern Biotech) were added. A fluorescence microscope (LSM700; Zeiss, Oberkochen, Germany) was used for taking the pictures. All pictures were processed with the ZenBlue 2012 Software Version 1.1.2.0. (Carl Zeiss Microscopy GmbH, Jena, Germany).

### Albumin endocytosis assay

UdRPCs were plated at a density of 40% without coating. After two days the cells were washed 1X with PBS and incubated in new medium contained 20 μg/ml of bovine serum albumin (BSA)-Alexa Fluor 488 conjugate (catalog no. A13100; Thermo Fischer) for 1 h at 37°C. Thereafter, the cells were washed three times with ice-cold PBS and fixed with 4% PFA for 15 min. Cell-associated fluorescence was analyzed using an excitation wavelength of 488 nm and an emission wavelength of 540 nm and imaged using a florescence microscope (LSM700; Zeiss, Oberkochen, Germany).

### Western blot analysis

For protein extraction, cells were harvested and lysed in RIPA buffer (Sigma Aldrich) supplemented with complete protease and phosphatase inhibitors cocktail (Roche). The lysates were separated on a 4-20% Bis-Tris gel and blotted onto a 0.45 µm nitrocellulose membrane (GE Healthcare Life Sciences). The membranes were then blocked with 5% respective primary antibodies: Total Smad 1 (CST, 1:1000, TBS-T 5% BSA), phospho Smad 1/5/8 (CST, 1:1000, TBS-T 5% milk), Total Smad 2/3 (CST, 1:1000, TBS-T 5% BSA), and phospho Smad 2/3 (CST, 1:1000, TBS-T 5% milk). After incubation with the appropriate secondary antibodies, signals were acquired with a Fusion-FX7 imaging system.

### Analysis of cell proliferation

Cell proliferation were analysed using resazurin metabolic colorimetric assay. UdRPCs were seeded (1×10^4^ cells/well) in a 6-well plate and incubated at 37°C in a humidified atmosphere at 5% CO_2_. The medium was substituted with 10% of a resazurin solution (0.1 mg/ml resazurin salt solution (Sigma-Aldrich) in PBS) with an end-volume of 2 ml per well and changed on daily basis. The cultures were incubated for 4h at 37°C in 5% CO_2_. Following this incubation period, the resazurin-containing medium was collected and the rate of resazurin conversion to resofurin by metabolically active cells was evaluated by spectrophotometric analysis at 570 and 600 nm. A final optical density (O.D.f) was determined for each sample, as follows: (O.D. 570/O.D.600)-(O.D.570c/O.D. c 600), where ‘OD.c’ are the O.D.s of control samples (fresh medium supplemented with resazurin, never in contact with cells). This procedure was carried out for 9 days, at the same hour, in triplicate.

### Cell lines used in this study and culture condition

The fibroblast cell used in this study were obtained from human foreskin fibroblast (HFF1) (ATCC, #ATCC-SCRC-1041, Manassas, VA, USA, www.atcc.org). Pluripotent stem cells (HFF-iPSCs (human foreskin fibroblast-derived induced pluripotent stem cells (iPSCs)) and ESCs (H1 (#WA01) and H9 (#WA09), WiCell Research Institute, Madison, WI, USA, www.wicell.org) were cultured in mTeSR on cell culture dishes coated with Matrigel (BD). Media were replaced change every day. Passaging of pluripotent stem cells was carried out with a splitting ratio of 1:3 to 1:10. Passaging was conducted manually using a syringe needle and a pipette under a binocular microscope or using a cell scraper and PBS (––).

