## Supplemental tables and figures for "A Comprehensive Molecular Portrait of Human Urine-derived Renal Progenitor Cells"

**Supplemental Material Table of Contents**

| **Supplemental Number** | **Legends** |
| --- | --- |
| **Supplemental Table S1** | List of Antibodies used for immunocytochemistry/flow-cytometry. |
| **Supplemental Table S2** | List of primers |
| **Supplemental Table S3 (Excel file)** | Sheet 1: Subsets of venn diagram comparison of UdRPCs of sample UM51 vs. hREPCs.  Sheet 2: Overrepresented GOs in the exclusive UM51 subset with 566 genes from the venn diagram comparison of UdRPCs of sample UM51 vs. hREPCs.  Sheet 3: Overrepresented GOs in the exclusive hREPCs subset with 438 genes from the venn diagram comparison of UdRPCs of sample UM51 vs. hREPCs.  Sheet 4: Overrepresented GOs in the up-regulated genes (limma-p-value < 0.05, ratio > 2) from the overlap subset of the venn diagram comparison of UdRPCs of sample UM51 vs. hREPCs.  Sheet 5: Overrepresented GOs in the down-regulated genes (limma-p-value < 0.05, ratio < 0.5) from the overlap subset of the venn diagram comparison of UdRPCs of sample UM51 vs. hREPCs |
| **Supplemental Table S4 (Excel file)** | Sheet 1: Subsets of venn diagram comparison of UdRPCs vs. fibroblasts.  Sheet 2: Overrepresented GOs in the exclusive UdRPC subset with 463 genes from the venn diagram comparison of UdRPCs vs. fibroblasts |
| **Supplemental Table S5 (Excel file)** | Sheet 1: Subsets of venn diagram comparison of iPSCs derived from UdRPCs (UdRPC_iPSCs), iPSCs derived from human foreskin fibroblasts (B4_HFF_iPSCs) and human embryonic stem cells (ESCs).  Sheet 2: Overrepresented GOs in the exclusive UdRPC_iPSCs subset with 150 genes from the venn diagram comparison of UdRPC_iPSCs, B4_HFF_iPSCs and ESCs.  Sheet 3: Overrepresented GOs in the exclusive B4_HFF_iPSCs subset with 312 genes from the venn diagram comparison of UdRPC_iPSCs, B4_HFF_iPSCs and ESCs.  Sheet 4: Overrepresented GOs in the exclusive ESCs subset with 197 genes from the venn diagram comparison of UdRPC_iPSCs, B4_HFF_iPSCs and ESCs. |
| **Supplemental Table S6 (Excel file)** | Sheet 1: Subsets of venn diagram comparison of UdRPCs treated with CHIR99021 vs. untreated UdRPCs.  Sheet 2: The set of 2491 up-regulated genes (p<0.05, ratio>1.33) from the venn diagram intersection of UdRPCs treated with CHIR99021 vs. untreated UdRPCs.  Sheet 3: The set of 2043 down-regulated genes (p<0.05, ratio<0.75) from the venn diagram intersection of UdRPCs treated with CHIR99021 vs. untreated UdRPCs.  Sheet 4: The set of 7255 not regulated genes (p>0.05, 0.75<ratio<1.33) from the venn diagram intersection of UdRPCs treated with CHIR99021 vs. untreated UdRPCs.  Sheet 5: Overrepresented KEGG pathways in the set of 2491 up-regulated genes from the venn diagram intersection of UdRPCs treated with CHIR99021 vs. untreated UdRPCs.  Sheet 6: Overrepresented KEGG pathways in the set of 2043 down-regulated genes from the venn diagram intersection of UdRPCs treated with CHIR99021 vs. untreated UdRPCs.  Sheet 7: Novel genes beginning with LOC (without published symbol) in the set of 2491 up-regulated genes (p<0.05, ratio>1.33) from the venn diagram intersection of UdRPCs treated with CHIR99021 vs. untreated UdRPCs.  Sheet 8: Novel genes beginning with LOC (without published symbol) in the set of 2043 down-regulated genes (p<0.05, ratio<0.75) from the venn diagram intersection of UdRPCs treated with CHIR99021 vs. untreated UdRPCs.  Sheet 9: Novel genes beginning with LOC (without published symbol) in the set of 7255 down-regulated genes (p>0.05, 0.75<ratio<1.33) from the venn diagram intersection of UdRPCs treated with CHIR99021 vs. untreated UdRPCs. |

**Supplemental Material Table of Contents**

| **Supplemental Number** | **Legends** |
| --- | --- |
| **Supplemental Figure S1** | Detailed analysis by bisulfite sequencing of CpG island methylation patterns within the 5´- regulatory region of the OCT4 gene in UM51 (control) and its iPSC derivative. Detailed CpG methylation profiles of the OCT4 5´-regulatory region are documented as revealed by bisulfite sequencing. Filled circles (black) denote methylated CpG dinucleotides, white denote unmethylated CpGs and gray CpG dinucleotides of unknown methylation status. Arrows indicate the transcription start site. |
| **Supplemental Figure S2** | (A) Immuno-phenotyping for MSC markers. (B) Secretome profile membrane. (C) Secretome genes related GOs and KEGG Pathways. (D) PODXL and CK19 staining. |
| **Supplemental Figure S3** | **Generation and characterization of iPSCs from UdRPCs.** |
| **Supplemental Figure S3_A** | (A) The reprogrammed vector-free UdRPCs-iPSCs stained positive for the transcription factors Oct4, Sox2 and Nanog and for the surface markers TRA-1-60, TRA-1-81 and SSEA-4 confirmed by immunofluorescence. |
| **Supplemental Figure S3_B** | (B) The immunofluorescence-based study shows a successful undirected differentiation into the mesoderm lineage detected with the mesoderm marker α-SMA and a successful specification along the endoderm layer confirmed with AFP, as well as the ectoderm layer proved with Nestin. |
| **Supplemental Figure S3_C** | (C) Dendrogram resulting from hierarchical clustering of global gene expression profiles of UdRPCs-iPSCs, UdRPCs, and established ESCs (H1, H9). Transcriptomes of UdRPCs-iPSCs cluster with H1, H9 while those of the UdRPCs cluster separately. Pearson correlation analysis of transcriptome data revealed a high correlation (green) of UdRPCs-iPSCs with ESCs but low correlation with UdRPCs. Pearson's correlation coefficient was calculated in which each replicate was pairwise compared with each other replicate. A value of 1 indicates perfect linear correlation while a value of 0 implies no correlation. |
| **Supplemental Figure S3_D** | (D) The origin of the formed iPSCs was assigned to its donor UdRPC line by determining the individual DNA signature. For this purpose, a PCR-based DNA fingerprinting using specific primer sets, which amplify different VNTRs (variable number of tandem repeats) was employed. Ultimately, the genotyping provide evidence, that reprogrammed iPSC clones originate from their parental UdRPC line and hence exclude the possibility of cross-contamination. Collecting all data, the excellent quality and integrity of the reprogrammed urinary progenitor cells was proven by a normal 46, XY karyotype. |
| **Supplemental Figure S4** | ESC-exclusively GOs when compared to HFF-iPSCs and UdPC-iPSCs. Treemap summarizing the GO-BP terms overrepresented in the 197 genes expressed exclusively in ESCs. The largest most significant group is associated with “negative regulation of astrocyte differentiation”, second comes “anion transmembrane transport”. |
| **Supplemental Figure S5** | KEGG pathways associated with genes up and down regulated upon CHIR treatment |

**Supplemental Table S1.** List of Antibodies used for immunocytochemistry/flow-cytometry.

|  | **Antibody** | **Dilution** |  |
| --- | --- | --- | --- |
| Renal progenitor marker | SIX2 | 1:200 | Proteintech Group Cat# 11562–1-AP, RRID:AB_2189084 |
| Pluripotency markers | CD133 | 1:500 | US Biological Cat# C251490B, RRID:AB_2284567Boster Biological; PA2049  |
| Pluripotency markers | Rabbit anti-OCT4 | 1:400 | Cell Signaling Technology Cat# 2840S, RRID:AB_2167691 |
| Pluripotency markers | Rabbit anti-SOX2 | 1:400 | Cell Signaling Technology Cat# 3579S, RRID:[AB_2195767](http://antibodyregistry.org/AB_2195767) |
| Pluripotency markers | Rabbit anti-NANOG | 1:800 | Cell Signaling Technology Cat# 4903S, RRID:AB_10559205 |
| Pluripotency markers | Mouse anti-TRA-1-60 | 1:1000 | Cell Signaling Technology Cat# 4746S, RRID:AB_2119059 |
| Pluripotency markers | Mouse anti-TRA-1-81 | 1:1000 | Cell Signaling Technology Cat# 4745S, RRID:AB_2119060 |
| Pluripotency markers | Mouse anti-SSEA4 | 1:1000 | Cell Signaling Technology Cat# 4755S, RRID:[AB_1264259](http://antibodyregistry.org/AB_1264259) |
| Differentiation markers | Mouse anti-Sox17 | 1:50 | R and D Systems Cat# AF1924, RRID:[AB_355060](http://antibodyregistry.org/AB_355060) |
| Differentiation markers | Rabbit anti-AFP | 1:200 | Cell Signaling Technology Cat# 2137S, RRID:AB_2209744 |
| Differentiation markers | anti-Nestin | 1:250 | Sigma-Aldrich Cat# N5413, RRID:AB_1841032 |
| Differentiation markers | anti-aSMA | 1:1000 | Dako Cat# M0851, RRID:AB_2223500 |
| MSC Marker | Vimentin (5G3F10) mouse mAb | 1:200 | Cell Signaling Technology, USA, 3390 |
| Renal Marker | Rabbit anti-CK19, | 1:100 | Novus Biologicals, NB100-687 |
| Pluripotency markers | C-Kit (H-300) rabbit polyclonal IgG | 1:200 | Tebu Bio, Germany |
| Renal Marker | CITED1, | 1:160 | Invitrogen, PA5-40585 |
| Renal Marker | PAX8 (D2S2I) rabbit mAb, | 1:200 | Cell Signaling Technology, USA, 59019s |
| Renal Marker | rb Anti WT1 clone 6F-H2, | 1:200 | MD Millipore Corp., O5-753 |
| Renal Marker | rb Nephrin AB, | 1:200 | Invitrogen, PA5-20330 |
| Renal Marker | rb PODXL, | 1:200 | Santa Cruz Biotech. sc33138 |
| Renal Marker | rb LIM1/LHX1, | 1:500 | Abcam, ab14554 |

**Supplemental Table S2.** List of primers

| Genes Name | Primer Sequences | Product  size (bp) |
| --- | --- | --- |
| BMP7 | F1: CAACCTCGTGGAACATGACAAG, R1: AAGATCAAACCGGAACTCTCGAT | 70 |
| CD133 | F1: GACTTGCGAACTCTCTTGAATGA, R1: GGTAGTGTTGTACTGGGCCAAT | 222 |
| RPL37A | F1: GTGGTTCCTGCATGAAGACAGTG, R1: TTCTGATGGCGGACTTTACCG | 84 |
| OCT4 | F1: GAGGGAGAGAGGGGTTGAGTAGTTTT,R1:ACTCCAACTTCTCCTTCTCCAACTTC | 469 |

**Supplemental Figure S1**

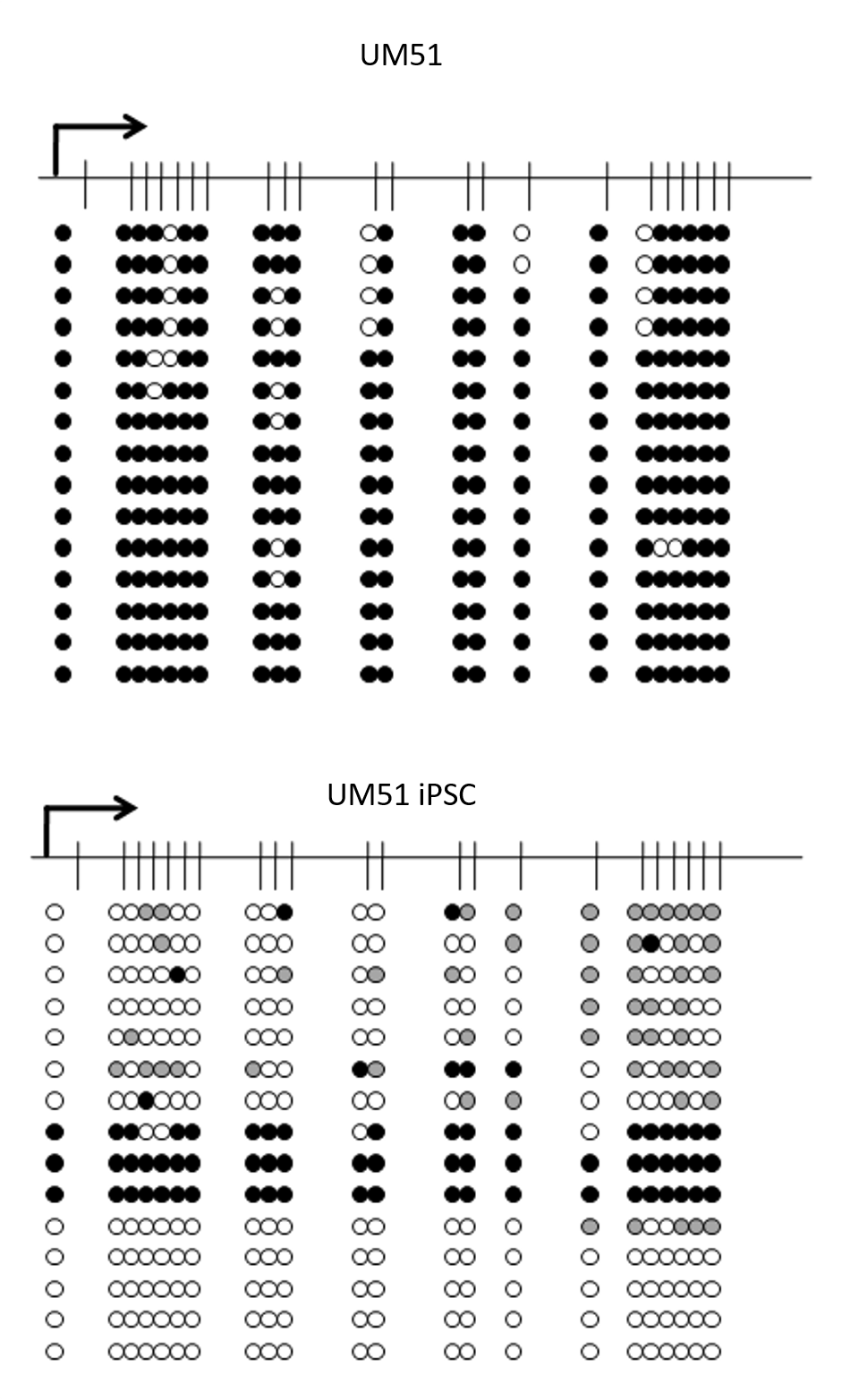

**Supplemental Figure S2**

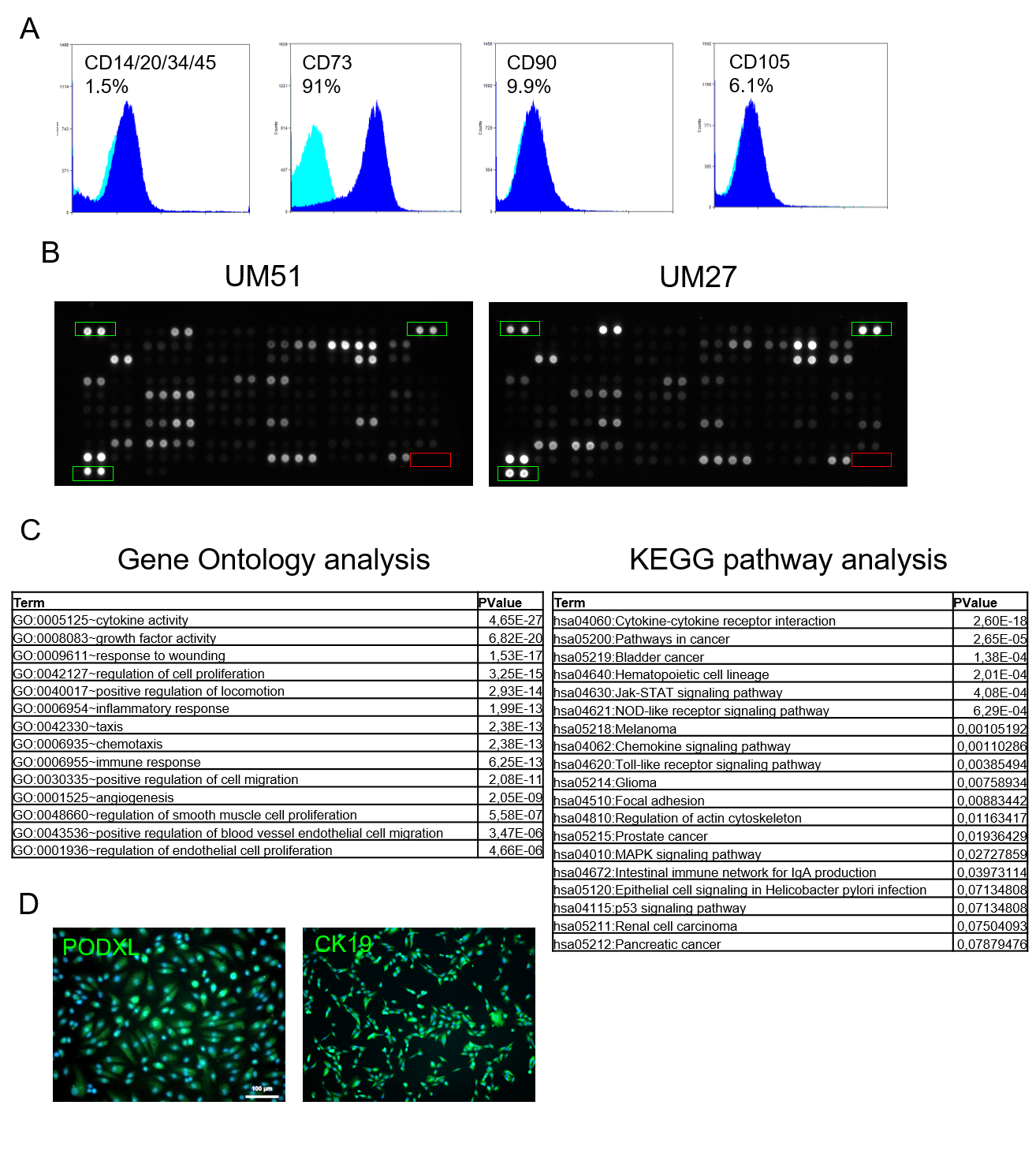

**Supplemental Figure S3**

**Supplemental Figure S3 A**

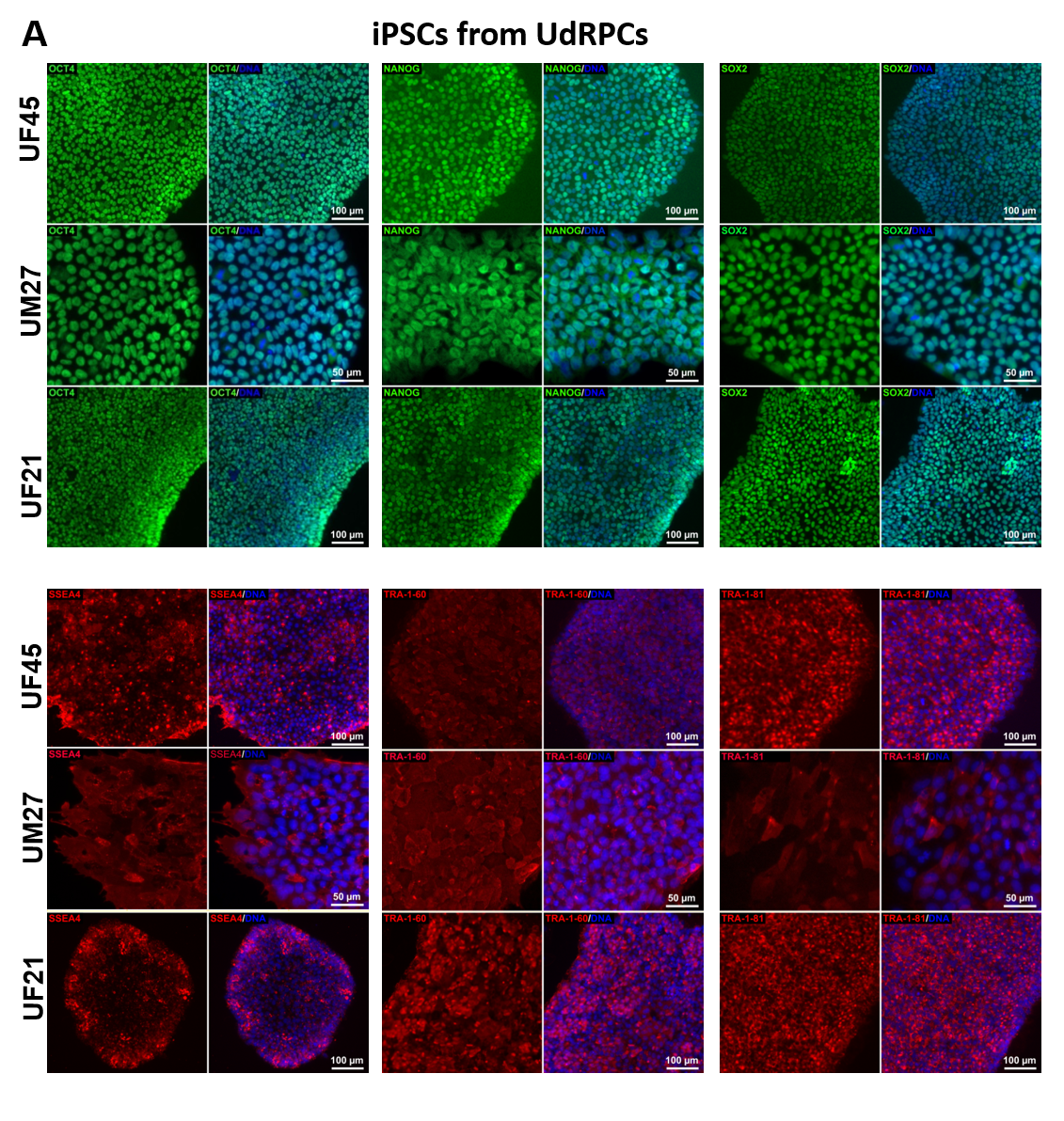

**Supplemental Figure S3 B**

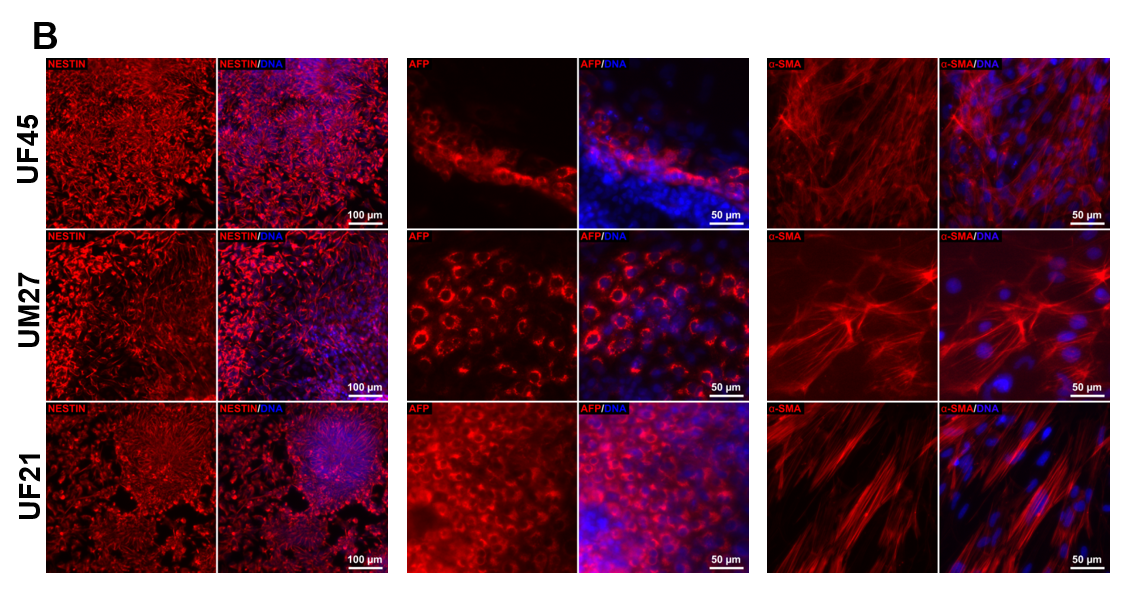

**Supplemental Figure S3 C**

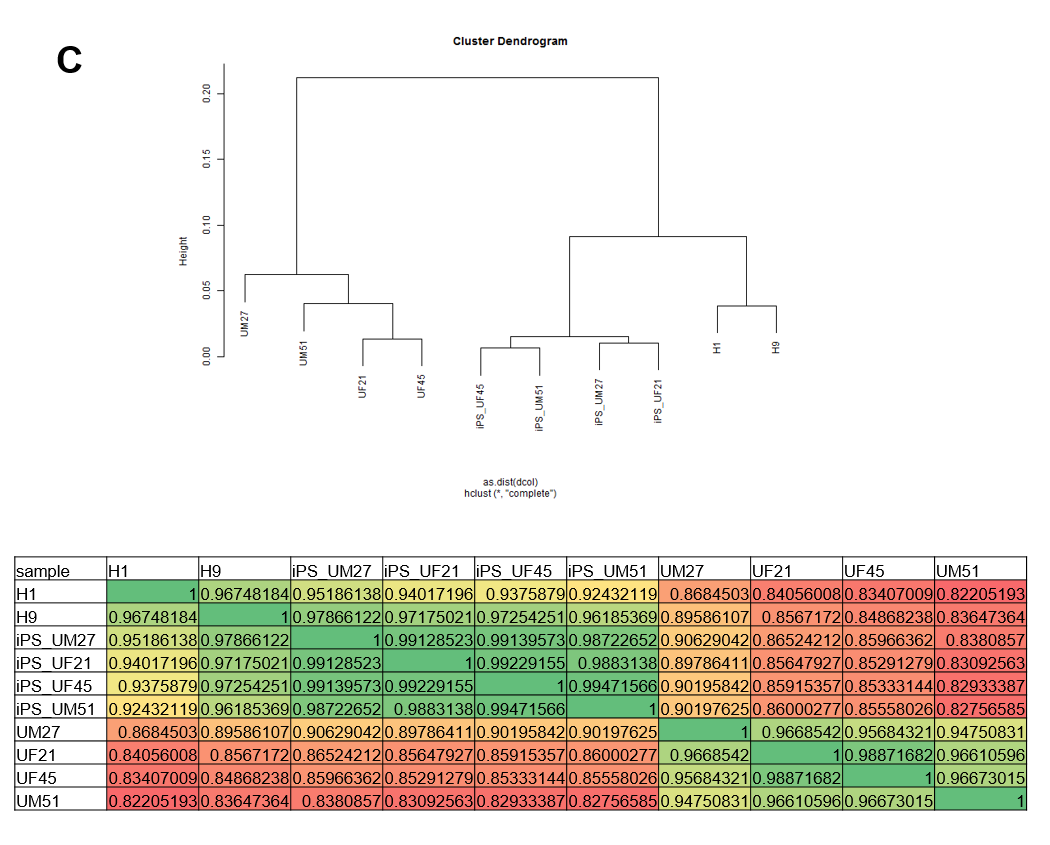

**Supplemental Figure S3 D**

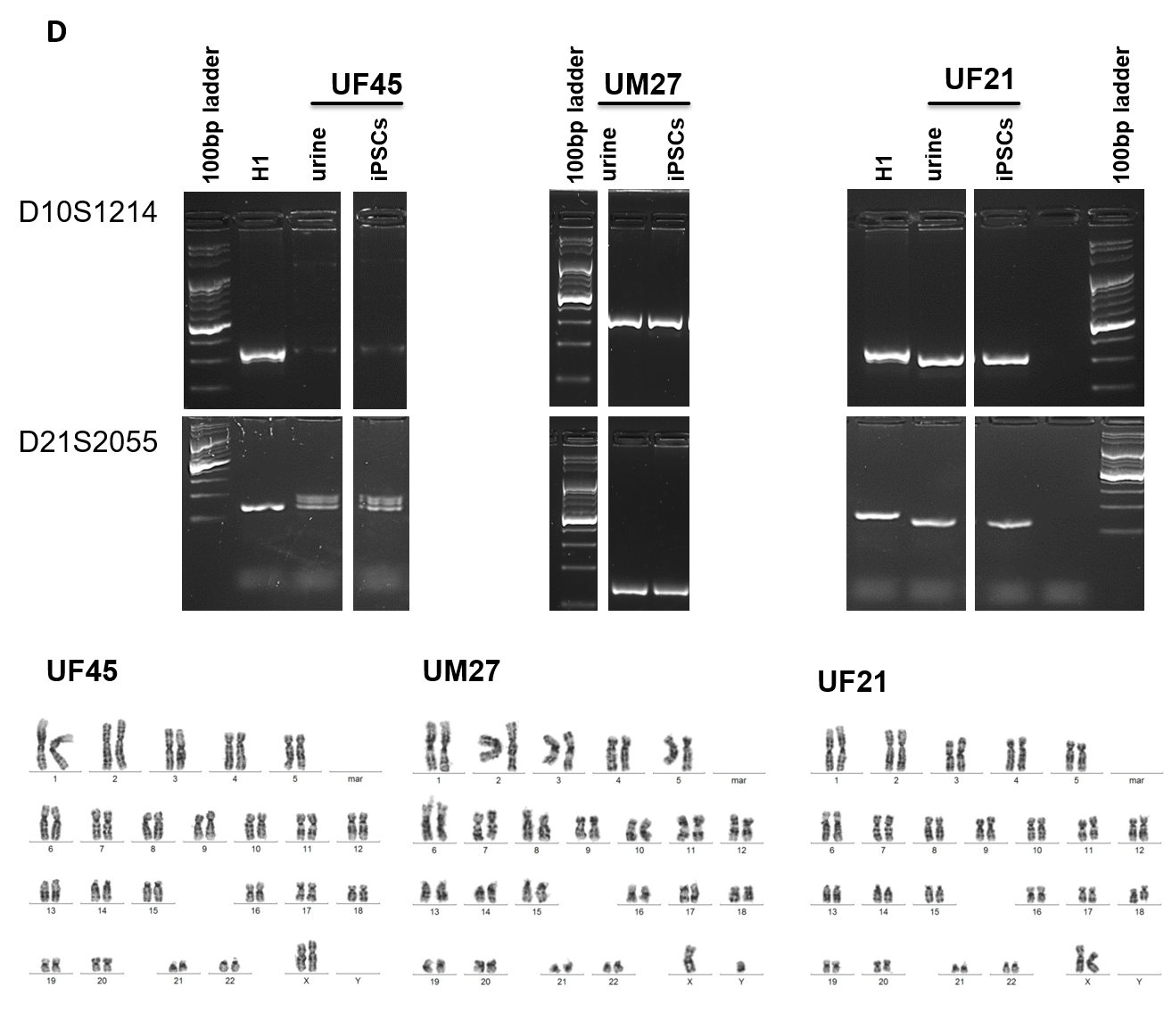

**Supplemental Figure S4**

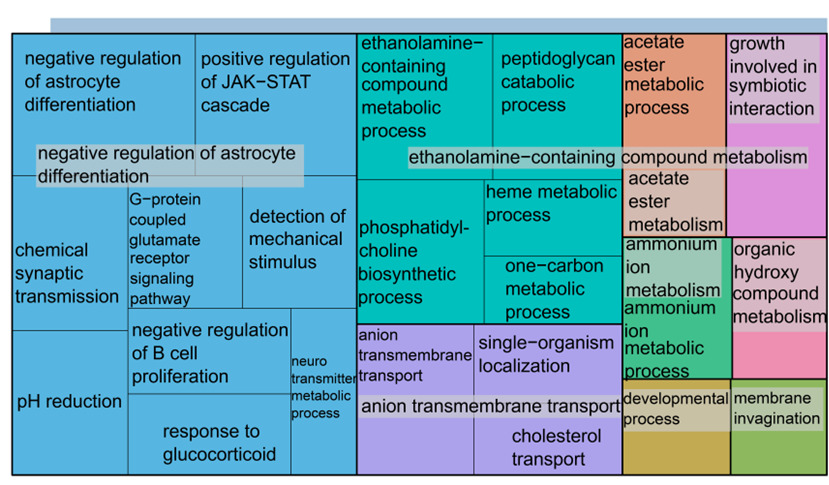

**Supplemental Figure S5**

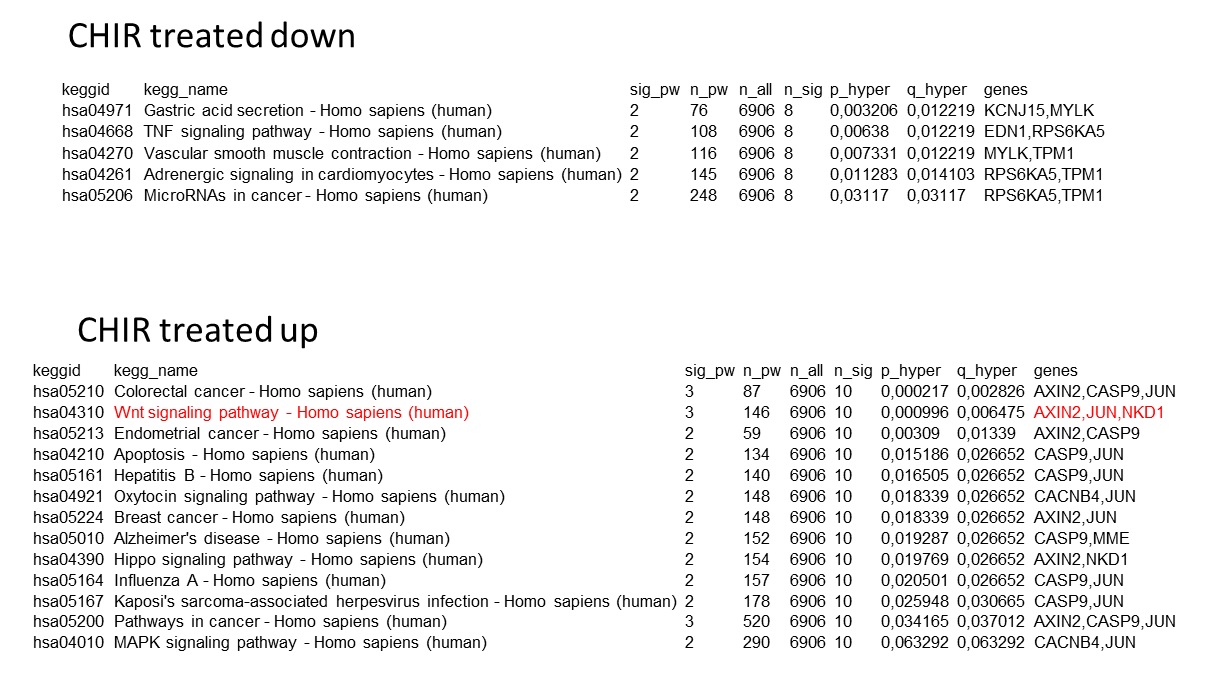
